# Compartmentalization of mRNAs in the giant, unicellular green algae *Acetabularia acetabulum*

**DOI:** 10.1101/2020.09.18.303206

**Authors:** Ina J. Andresen, Russell J. S. Orr, Kamran Shalchian-Tabrizi, Jon Bråte

**Affiliations:** Section for Genetics and Evolutionary Biology, Department of Biosciences, University of Oslo, Kristine Bonnevies Hus, Blindernveien 31, 0316 Oslo, Norway; Natural History Museum, University of Oslo, Oslo, Norway; Centre for Epigenetics, Development and Evolution, Department of Biosciences, University of Oslo, Kristine Bonnevies Hus, Blindernveien 31, 0316 Oslo, Norway

**Keywords:** Acetabularia acetabulum, Dasycladales, UMI, STL, compartmentalization, single-cell, mRNA

## Abstract

*Acetabularia acetabulum* is a single-celled green alga previously used as a model species for studying the role of the nucleus in cell development and morphogenesis. The highly elongated cell, which stretches several centimeters, harbors a single nucleus located in the basal end. Although *A. acetabulum* historically has been an important model in cell biology, almost nothing is known about its gene content, or how gene products are distributed in the cell. To study the composition and distribution of mRNAs in *A. acetabulum*, we have used quantitative RNA-seq to sequence the mRNA content of four sections of adult *A. acetabulum* cells. We found that although mRNAs are present throughout the cell, there are large pools of distinct mRNAs localized to the different subcellular sections. Conversely, we also find that gene transcripts related to intracellular transport are evenly distributed throughout the cell. This distribution hints at post-transcriptional regulation and selective transport of mRNAs as mechanisms to achieve mRNA localization in *A. acetabulum*.

## Introduction

Large and complex morphological forms are predominantly found among multicellular organisms such as animals, plants and kelps. However, also several single celled organisms have elaborate cellular morphologies, and it has therefore been previously argued that multicellularity is not a requirement for the formation of structural complexity (Kaplan, 1992; Kaplan et al., 1991; Niklas et al., 2013; Ranjan et al., 2015). By subcellular compartmentalization of RNA or proteins, single-celled organisms can chamber their bodies into differently shaped subunits, further facilitating development of sophisticated body plans without cellularization (Kaplan, 1992; Kaplan et al., 1991; Niklas et al., 2013). The green algae order Dasycladales harbors multiple examples of single-celled species with highly elaborate and complex cellular morphologies. With only a single nucleus, these algae have evolved into numerous morphological forms, and can grow up to 20 cm long (Berger, 1990, 2006).

Being used as a model organism to study the link between genetics and cellular development, *Acetabularia acetabulum* is the most studied dasycladalean species. This alga is a tropical, marine species found in shallow waters in the Mediterranean Sea, Northern Africa and South-West Asia. Divers often refer to *A. acetabulum* as “mermaids wineglass” because of its distinctive morphology; the basal rhizoid, which hosts the nucleus as well as ensuring anchoring of the cell to a substrate, is followed by a naked, elongated stalk with a slightly concave and disc-looking structure (cap), at the apical end (Figure 1A).

**Figure 1.**
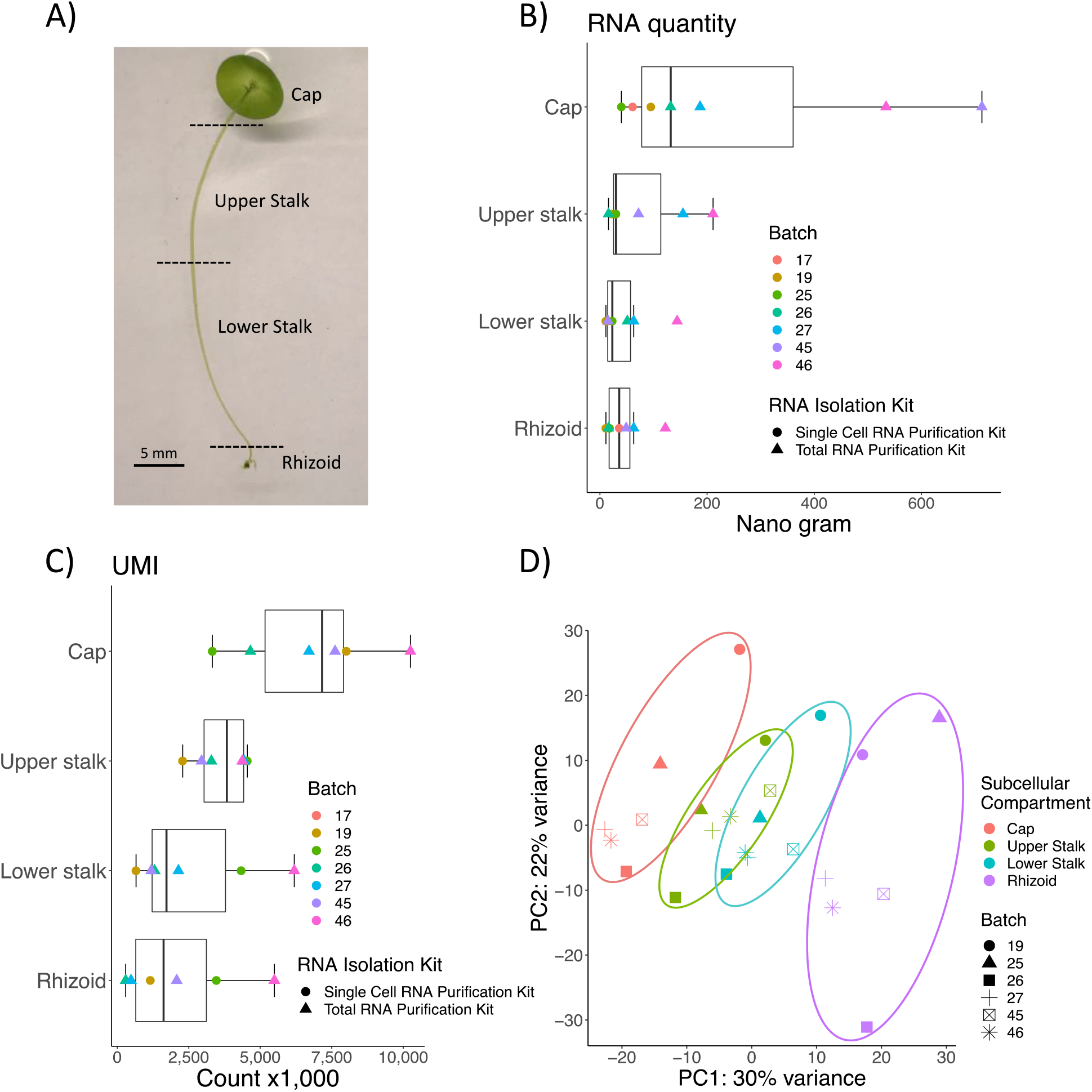
RNA isolation and sequencing of subcellular compartments of adult *A. acetabulum*. A) Image of an adult cell of *A. acetabulum*. The dashed lines indicate approximate incision sites for separating the different subcellular sections; cap, upper stalk, lower stalk and rhizoid. B) Boxplot showing the total RNA quantity isolated from the different subcellular compartments. The dots represent the individual samples colored according to which batch the sample originate from, and shaped according to which to which RNA isolation kit that was used. C) Boxplot showing the summarized gene expression levels (total UMI counts) of the different samples. The dots are the same as above. D) Principal Component Analysis (PCA) of the sample variation based on variance stabilized counts (see Methods). The four subcellular compartments are shown in color, and the different batches which the samples originate from (described in the Methods) are indicated as shapes.

Through his ground breaking amputation and grafting experiments, Joachim Hämmerling was the first to identify the presence of substances controlling the subcellular morphogenesis of *A. acetabulum* (Hämmerling, 1934a, 1934b; Hämmerling, 1953). He also showed that these substances were distributed in gradients, with the highest accumulation at the cap and rhizoid regions (Hämmerling, 1934c; Hämmerling, 1963). These morphogenetic substances were later shown to consist of mRNA (Baltus et al., 1968; Garcia et al., 1986; Kloppstech, 1977; Naumova et al., 1976), but the content of the RNA gradient, and whether it was composed of homogeneously or differentially distributed RNA transcripts was not known.

A few studies have tried to decipher the RNA gradient in *A. acetabulum*. Naumova et al. (1976) argued that the gradient was due to differential metabolism of chloroplast ribosomal RNA along the cell rather than differential transportation of nuclear mRNA. A more recent study by Vogel et al. (2002) examined the expression of 13 house-keeping genes and found differential accumulation of several gene transcripts, where some transcripts seemed to accumulate in the rhizoid, some in the cap, while others were evenly distributed along the cellular body. Serikawa et al. (1999) demonstrated that the *A. acetabulum*-specific homeobox-containing gene, *Aaknox1*, shifted localization from being evenly distributed during vegetative growth, to basal accumulation in the final stages of the *A. acetabulum* life cycle. And both the transcripts encoding carbonic anhydrases as well as their translated protein products were shown to be co-localized in the apical parts of adult *A. acetabulum* cells (Serikawa et al., 2001), indicating that localization of mRNA is a mechanism for correct localization of protein. Altogether, these studies suggested that subcellular localization of mRNA is common in the *A. acetabulum* cell, and an important mechanism for establishment of subcellular structures.

mRNA localization has also been demonstrated in another gigantic single-celled green algal genus, *Caulerpa* (Arimoto et al., 2019; Ranjan et al., 2015). These single-celled organisms contain hundreds of nuclei distributed across the cell and the localization of mRNA is achieved, at least in part, by differential nuclear transcriptional regulation in the different parts of the cell (Arimoto et al., 2019). Unlike *Caulerpa, A. acetabulum* has only a single nucleus (located in the basal rhizoid) and clearly the mechanisms behind the subcellular localization of mRNAs, and ultimately the establishment of the cellular body plan, must be different. Although these mechanisms are unknown, there are several indications that mRNAs are actively transported along the cytoskeleton in *A. acetabulum*. Kloppstech et al., (1975a and 1975b) showed that ribosomal- and polyadenylated RNA travel with different speeds in *A. acetabulum*, and that polyadenylated RNA moved with a speed of 0.2 μm/s, which is much faster than movement with normal diffusion, suggesting that RNA transportation is both active and specific. Further, staining experiments performed on *A. peniculus* showed that actin proteins and polyadenylated RNAs were co-localized, and that treatment with cytochalasin D (inhibitor of actin polymerization) lead to a disruption of already established mRNA gradients (Mine et al., 2001). These findings supported the theory that the cytoskeleton is involved in polar transportation of mRNAs in *Acetabularia* species (Vogel et al., 2002).

The studies on the expression and localization of mRNAs in *A. acetabulum* has so far only been performed on a restricted number of genes, and it is not known how general this phenomenon is, and how gene localization is related to the observed gradients of RNA along the cell. We have therefore in this study characterized the expression profile of all mRNAs in adult *A. acetabulum* cells. To achieve this, we have exploited recent developments in single-cell RNA-seq technology which allowed us to quantitively sequence mRNAs from four subcellular sections of the cell, the upper and lower parts of the stalk, and the rhizoid (Figure 1A). We investigate whether subcellular compartmentalization is a general phenomenon for all types of mRNAs in *A. acetabulum*, and whether the distribution of mRNAs can be coupled to the formation of body plan and the complex cellular morphology of this algae.

## Methods

### Culturing *Acetabularia acetabulum* cells

*A. acetabulum* cells were cultured in cell/tissue culture flasks in Dasycladales Seawater Medium prepared after the recipe of UTEX Culture Collection of Algae at The University of Texas at Austin (https://utex.org/products/dasycladales-seawater-medium?variant=30991770976346). The medium was changed biweekly, and the algae were kept in incubators with a 12/12h light/dark cycle (light intensity of 2500 lux) with a constant temperature of 20°C.

### Dissection of cells and RNA isolation

The *A. acetabulum* cells were 5-8 cm long with fully grown caps, but no apparent gametes in the gametangia, and no whorls along the stalk (Figure 1A). The cells were washed three times in 1X PBS to remove residues from the medium, and dissected into four subcellular regions; the “cap” (incision about 2 mm from the apical tip or just below the cap), “rhizoid” (incision about 2 mm above the rhizoid), “upper stalk” (the upper half of the stalk) and “lower stalk” (the lower half of the stalk). Dissection was carried out in dry petri dishes to limit cytoplasmic loss, and new, sterile scalpels were used between incision. Subcellular regions from 5 – 8 adult cells were pooled together to achieve sufficient RNA quantities. The procedure was repeated to create seven replicates of RNA extraction. The subcellular samples are numbered according to which batch of cells they originate from, e.g. the samples named “Cap 19” and “Rhizoid 19” are the cap- and rhizoid sections from the same batch of individuals (batch19).

The dissected pieces were transferred to green MagNA Lyser Green Beads (Roche Life Science, Germany), containing lysis buffer (see below), and flash frozen in liquid nitrogen. RNA from 3 batches (batch 17, 19 and 25) was isolated using the “Single Cell RNA Purification Kit” (NORGEN BIOTEK CORP, Canada) with an 8 ul elution buffer. RNA from the remaining 4 batches (batch 26, 27, 45 and 46) was isolated using the “Total RNA Purification Kit” (NORGEN BIOTEK CORP, Canada) with 40 ul elution. RNA quality and quantity were inspected on the Agilent 2100 Bioanalyzer using the Agilent RNA 6000 Pico kit (Agilent Technologies, Inc, Germany).

### Library preparation and sequencing

ERCC RNA Spike-In Mix I (ThermoFisher Scientific, Massachusetts, USA) was added to each sample before mRNA enrichment. The amount of added ERCC Spike-In Mix was adjusted to the amount of RNA in each sample, according to the manufacture’s protocol. mRNA enrichment was performed using NEXTflex™ Poly(A) Beads (BIOO Scientific Corporation, Texas, USA) before library preparation with the NEXTflex Rapid Directional qRNA-Seq Library Prep kit for Illumina sequencing (BIOO Scientific Corporation, Texas, USA). This library preparation kit assigns a unique molecular identifier (UMI), or Stochastic Label (STL), to both ends of the mRNA fragments after enzymatic fragmentation, but before cDNA synthesis and amplification. This allows for distinguishing between PCR duplicates and true identical sequences which map to the same loci, ensuring a better quantitative representation of the original number of mRNA fragments in the samples than standard RNA-seq library protocols without UMI-labelling (Toloue et al., 2013). A total of 30 PCR cycles were run for sample 17.1-4, 25 cycles were run for sample 25.4 and 26.4, and 20 cycles were run for sample 19.1-4, 25.1-3, 26.1-3, 27.1-4, 45.1-4 and 46.1-4 to create libraries of approximately equal quantities as measured by gel electrophoresis.

The 28 libraries were sequenced on the Illumina HiSeq4000 platform producing 150 bp Paired-End sequences (with an insert size of 350 bp). The sequencing was performed at the Norwegian Sequencing Centre (www.sequencing.uio.no) at the University of Oslo.

### *De novo* transcriptome assembly and annotation

In order to obtain a complete transcriptome representing as many transcripts as possible to use for gene quantification, the resulting sequences from the 28 adult sequence libraries were assembled together with 20 transcriptome sequence libraries from various developmental stages of *A. acetabulum* (unpublished data generated by our research group). Briefly, the assembly was performed by first removing the nine first UMI bases of each sequence (from the 5’ end) with Trimmomatic v/0.35 (Bolger et al., 2014). Further, the 3’-adaptor sequences, and low-quality sequences (phred score < 20) were trimmed. Only sequences longer than 36 bp were retained. An additional trimming with TrimGalore v/0.3.3 (http://www.bioinformatics.babraham.ac.uk/projects/trim_galore/) was performed to remove any remaining adaptor sequences. ERCC RNA spike-ins were removed from the data set by mapping the reads to the known ERCC RNA spike-in sequences using Bowtie 2 v/2.2.9 (Langmead et al., 2012). A total of 2,095,599,508 paired end reads (1,047,799,754 pairs) were obtained after preprocessing and used for transcriptome assembly. *De novo* assembly was performed using Trinity v/2.5.1 (Grabherr et al., 2011). To reduce the number of possible mapping sites in downstream analysis, the transcriptome was reduced to the highest expressed isoform for each gene. These isoforms were found by subsampling 10% of the sequenced reads of every sample using the BBMap package (Bushnell, 2015), mapping them to the trinity assembly, and further extracting the isoforms with the highest coverage. Transcripts encoded by the chloroplast- and mitochondrial genome were identified by Megablast against a chloroplast database containing chloroplast genomes from 59 published green algae species, and a database containing mitochondrial genomes from 24 published green algae species (Table S1 and S2). Transcripts giving significant hits against the databases where further examined by Megablast against the NCBI Nucleotide collection database in order to exclude possible prokaryote contaminantion. Transcripts with no hits to either the plastid and mitochondrial databases were considered as nuclear encoded. As mRNA enrichment with poly(A) beads does not remove rRNA completely, rRNA transcripts were identified by Megablast against complete or partial 18S, 28S and 5.8S sequences from 20 green algae species (Table S3).

Transcriptome completeness was assessed by BUSCO v3.0 (Simao et al., 2015) analysis against the Chlorophyta and Eukaryote datasets. Since nuclear genes of *A. acetabulum* have an alternative codon usage, where TGA is the only stop codon, and TAA and TAG instead encodes glutamine (Jukes, 1996; Schneider et al., 1989), the alternative genetic code (translation table 6: Ciliate, Dasycladaen and Hexamita Nucelar Code) was used during BUSCO evaluation.

TransDecoder v/3.0.0 (Haas et al., 2013) was used to predict coding regions. Translation table 6 was used to translate nuclear encoded transcripts, translation table 16 (Chlorophycean Mitochondrial Code) was used to translate mitochondrial encoded transcripts and translation table 1 (Universal Code) was used to translate chloroplast encoded transcripts. Translation table 1 was also used for transcripts matching both the chloroplast and the mitochondrial database. The minimum peptide length was set to 70 amino acids, and the single best ORF per transcript Predicted peptide sequences were further annotated with the eggNOG-mapper v/5.0 (Huerta-Cepas et al., 2017; Huerta-Cepas et al., 2019).

**Table 1.** Total RNA isolation and mRNA sequencing of subcellular fragments of *Acetabularia acetabulum*. The naming of samples is described in the text. Read numbers are given as pairs of reads (single reads were discarded), both before and after trimming. Mapping rate describes the percentage of paired reads mapping concordantly (i.e. mapping in the expected orientation relative to each other) to the *de novo* assembled transcriptome, total UMI count is the sum of transcript expression levels for each sample after removing PCR duplicates. Expressed transcripts shows the number of transcripts with at least one UMI count. % of expressed transcripts with count ≤ 2 is the percentage of “low count” transcripts.

| Subcellular compartment | Batch | Total RNA (ng) | Raw reads (PE) | Trimmed reads (PE) | Mapped read pairs | Mapping rate (%) | Total UMI count | Expressed transcripts | % of transcripts with count $\leq 2$ |
| --- | --- | --- | --- | --- | --- | --- | --- | --- | --- |
| Cap | 17 | 61 | 50 770 695 | 5 820 120 | 3 585 755 | 75 | 584 | 290 | 79 |
| Upper stalk | 17 | 29 | 82 587 170 | 11 714 864 | 7 206 886 | 73 | 660 | 257 | 68 |
| Lower stalk | 17 | 14 | 2 092 | 260 | 166 | 74 | 0 | 0 | - |
| Rhizoid | 17 | 36 | 38 755 094 | 18 101 837 | 11 382 777 | 75 | 475 | 167 | 51 |
| Cap | 19 | 95 | 25 526 511 | 19 934 510 | 12 521 349 | 76 | 8 010 785 | 62 257 | 44 |
| Upper stalk | 19 | 22 | 14 007 423 | 10 572 421 | 6 395 304 | 73 | 2 285 994 | 45 187 | 42 |
| Lower stalk | 19 | 11 | 18 869 617 | 14 309 848 | 8 742 817 | 74 | 656 729 | 30 620 | 43 |
| Rhizoid | 19 | 11 | 24 207 912 | 18 101 837 | 11 021 291 | 74 | 1 156 418 | 38 275 | 45 |
| Cap | 25 | 40 | 22 444 960 | 19 460 102 | 11 548 241 | 72 | 3 325 240 | 58 967 | 45 |
| Upper stalk | 25 | 30 | 31 098 736 | 26 710 669 | 15 574 348 | 73 | 4 547 527 | 57 122 | 43 |
| Lower stalk | 25 | 23 | 37 346 096 | 32 191 425 | 19 013 248 | 73 | 4 334 098 | 62 677 | 44 |
| Rhizoid | 25 | 18 | 37 858 071 | 32 519 937 | 19 919 778 | 73 | 3 466 028 | 58 615 | 48 |
| Cap | 26 | 132 | 32 643 222 | 28 358 416 | 16 777 807 | 72 | 4 656 581 | 63 592 | 44 |
| Upper stalk | 26 | 16 | 22 694 589 | 19 609 743 | 11 646 681 | 72 | 3 291 813 | 50 613 | 42 |
| Lower stalk | 26 | 51 | 37 555 049 | 32 280 766 | 19 117 797 | 73 | 1 307 014 | 39 224 | 43 |
| Rhizoid | 26 | 16 | 27 307 714 | 16 273 913 | 9 951 940 | 72 | 293 461 | 23183 | 39 |
| Cap | 27 | 187 | 27 859 229 | 21 702 199 | 13 055 825 | 72 | 6 704 550 | 77 998 | 45 |
| Upper stalk | 27 | 155 | 30 596 065 | 23 362 341 | 14 172 132 | 73 | 4 437 570 | 59 649 | 42 |
| Lower stalk | 27 | 63 | 25 217 910 | 19 013 147 | 11 575 387 | 73 | 2 137 899 | 45 753 | 43 |
| Rhizoid | 27 | 63 | 29 781 682 | 15 189 655 | 9 203 797 | 73 | 480 492 | 33 149 | 46 |
| Cap | 45 | 713 | 30 795 845 | 26 732 918 | 16 179 143 | 73 | 7 615 615 | 71 764 | 45 |
| Upper stalk | 45 | 72 | 28 618 780 | 24 538 566 | 14 672 568 | 74 | 2 949 290 | 55 172 | 47 |
| Lower stalk | 45 | 15 | 29 851 320 | 25 462 402 | 15 279 659 | 73 | 1 186 788 | 36 048 | 43 |
| Rhizoid | 45 | 49 | 43 496 924 | 36 991 784 | 22 472 939 | 73 | 2 078 673 | 44 095 | 44 |
| Cap | 46 | 534 | 43 691 338 | 37 748 810 | 22 697 840 | 72 | 10 250 494 | 79 349 | 44 |
| Upper stalk | 46 | 211 | 26 586 549 | 22 842 662 | 13 930 603 | 73 | 4 370 405 | 61 734 | 45 |
| Lower stalk | 46 | 144 | 29 167 985 | 25 131 292 | 15 639 747 | 73 | 6 189 014 | 64 045 | 46 |
| Rhizoid | 46 | 122 | 28 000 697 | 24 124 084 | 14 737 583 | 72 | 5 492 434 | 75 135 | 48 |

### Gene expression quantification, normalization and sample clustering

In order to quantify the gene expression, processed reads (after removing UMI’s and low quality bases) were mapped to the transcriptome (the highest expressed isoform of each gene) with Bowtie2 v/2.2.9 (Langmead et al., 2012) and gene count files were generated using dqRNASeq (https://github.com/e-hutchins/dqRNASeq), a Unix script, developed for analyzing sequence data obtained with the NEXTflex Rapid Directional qRNA-Seq Library Prep kit. By using the resulting mapping file, together with the raw unprocessed reads, the script collapses paired reads with identical start- and stop sites *and* identical UMIs (assumed PCR duplicates) and count these as one, while fragments with identical start- and stop sites but different UMIs are counted individually (these are assumed to originate from different RNA fragments).

The plotCountDepth function in the SCnorm R package (Bacher et al., 2017) was used to calculate and visualize the relationship between sequencing depth and gene counts across samples. For the highest expressed genes (i.e. carrying the most robust signal) there was a positive relationship between sequencing depth and gene expression, and we therefore continued with the normalization procedure in the DESeq2 package (Love et al., 2014). To reduce the computational burden, we filtered the transcripts to have a minimum count of 1 (raw count) in at least 4 samples before DESeq2 normalization (these would not have affected the downstream statistical analysis as they would have been filtered out anyway).

To compare the similarity between samples based on the overall variation in transcript abundance, the normalized counts were transformed using the variance stabilizing transformation (VST) function in DESeq2 and the samples were clustered/visualized with PCA plots using the ggplot2 package (Wickham, 2016).

### Differential transcript abundance estimation

A Wald test (implemented in DESeq2), performed pairwise between the different subcellular compartments, was used to identify transcripts with differential abundance between at least two subcellular compartments. DESeq2 normalized read counts was used as input for the test. The batch origin of each sample was added as a blocking factor in the test (added to the design formula) to take into account any potential influence on the gene counts. Transcripts with adjusted p-value < 0.05 were considered as significantly differentially abundant (DE). Only nuclear encoded mRNAs were used in the DE test.

A Venn diagram was constructed using the systemPipeR package in R (Backman et al., 2016) to visualize and compare the DE transcripts of each subcellular compartment. To visualize and plot DE transcripts based on expression levels, the raw counts were converted to CPM’s (count per million) followed by TMM normalization (Trimmed Mean of M-values) using the edgeR package in R (McCarthy et al., 2012; Robinson et al., 2010). The mean expression values were further scaled and clustered in a heatmap using the Pheatmap package in R (https://CRAN.R-project.org/package=pheatmap).

### GO-enrichment analysis

GO-terms provided by EggNOG were converted to GO-slim with OmicsBox (https://www.biobam.com/omicsbox), and GO-enrichment analysis on the differentially distributed transcripts unique to the cap, upper stalk, lower stalk and rhizoid were performed using the R package GOseq (Young et al., 2010). In addition, the stalk samples were analyzed together by combining the uniquely differentially distributed transcripts from the upper- and lower stalk samples, as well as the differentially distributed transcripts shared between them. Lists of the unique transcripts from each subcellular compartment were extracted and inputted into GOseq, together with a list of transcript lengths, to account for any bias introduced from transcript length variation (longer transcripts might be easier to annotate or could receive more GO-annotations than shorter transcripts). GO-terms with a false discovery rate (FDR) < 0.05 were considered enriched. The “hit percentage” of each enriched GO-term within a subcellular compartment was calculated as the percentage of differentially distributed transcripts in a given GO-category compared to the number of transcripts in the transcriptome in the same GO-category. The hierarchical organization of the different enriched GO-terms were explored using the Mouse Genome Informatics web page (http://www.informatics.jax.org/vocab/gene_ontology). Heatmaps of GO terms showing the hit percentage of significantly enriched GOs was constructed using the ComplexHeatmap package in R (Gu et al., 2016), with a suitable number of K-means row-splitting. Row_km_repeats was set to 100, which runs clustering multiple times and outputs the consensus cluster.

### Annotation of transcripts related to intracellular transport and localization

Transcripts related to cytoskeletal components (Actin, Tubulin and related genes) and cytoskeletal motor proteins (Myosins, Dyneins and Kinesins), and poly(A) polymerases, were extracted from the eggNOG annotation. Homologs of genes related to cellular transport (COP and Clathrin) were identified by reciprocal blast by using annotated genes in NCBI RefSeq from *Chlamydomonas reinhardtii* as queries (see Table S4 for queries) against the *A. acetabulum* transcriptome (blastp value cutoff < 0.0001). The resulting hits of the *A. acetabulum* transcriptome were searched against Swissprot using blastp (evalue < 0.0001). Transcripts which did not produce hits against Swissprot of the same category as in the first blast search were discarded. Transcripts with a mean TMM-normalized CPM > 1 across all samples were plotted, with standard error, using the R package ggplot2.

### Comparative transcriptomics between *A. acetabulum* and *Caulerpa taxifolia*

The *Caulerpa taxifolia* transcriptome (Ranjan et al., 2015) was translated into amino acid sequences using Transdecoder v/3.0.0. Orthologous protein sequences between *A. acetabulum* and *C. taxifolia* were identified using Orthofinder v/2.3.3 (Emms et al., 2015). RSEM generated gene expression data from six different subcellular compartments of *C. taxifolia* (frond apex, pinnules, rachis, frond base, stolon and holdfast) was downloaded from the supplementary datafiles of Ranjan et al. (2015). The counts were rounded to the nearest integer and converted to TMM-normalized counts (as described above). The TMM counts from the single-copy orthologs from the different subcellular compartments of *A. acetabulum* and *C. taxifolia* were merged and the differences in sample variation was visualized using the prcomp function in R and the ggplot2 R package.

## Results

### Subcellular RNA isolation, sequencing and read processing

The highest amount of total RNA was extracted from the cap samples (average of 252 ng), followed by the upper stalk samples (average of 76 ng), the lower stalk (average of 46 ng), with the lowest amount extracted from the rhizoid samples (average of 45 ng) (Table 1 and Figure 1B. The highest yield of total RNA was obtained using the “Total RNA purification kit” with an elution volume of 40 ul (used for batch 26, 27, 45 and 46), which gave approximately four times as much total RNA compared to the “Single Cell RNA purification kit” with an elution volume of 10 ul (used for batch 17, 19 and 25) (Table S5). But still, the relative amounts of RNA isolated from the different samples were the same regardless of the isolation kit.

The sequence reads from batch 17 had overall very low-quality scores as well as a high number of duplicates (mostly from sequencing the adapters). Therefore, very few sequences were retained after filtering and almost no genes were detected in these samples. The samples from batch 17 were therefore discarded from further analyses. Between 14 and 43 million raw read pairs were produced from each of the remaining 24 samples. Quality trimming and removal of unpaired reads after trimming reduced the numbers by 13-25% for the majority of samples, except for the Rhizoid of batch 26 and 27, where trimming reduced the number of reads by 40 and 49% (Table 1). Still, more than 15 million read pairs were left for these samples.

### Transcriptome assembly and annotation

Assembling reads *de novo* produced an assembly consisting of 246,083 ‘genes’, or transcripts, with a total of 429,781 different isoforms (Table S6), where the longest transcript was 17,196 bp and the shortest 201 bp. 245,334 transcripts were considered nuclear encoded, 389 transcripts gave hits against the chloroplast database and were considered to be chloroplast encoded, 99 transcripts gave hits against the mitochondrial database and were considered mitochondrial encoded. 261 gave hits against both the chloroplast and mitochondrial databases and were considered as possible prokaryotic/unclassified transcripts as they also gave BLAST hits against prokaryotic sequences against NCBInr. 114,146 transcripts were predicted as protein coding. Of these, 113,900 belonged to the nuclear genes, 178 to the chloroplast, and 68 to the mitochondria. 38,131 transcripts gave ortholog hits when analyzed by EggNOG, of where 18 779 transcripts were assigned GO-terms.

Assessing the presence of conserved eukaryotic genes in the transcriptome with a BUSCO analysis estimated a ~96% completeness based on a pan-eukaryotic dataset and a ~70% completeness based on a Chlorophyta dataset (Table 2). The pan-eukaryote dataset is much smaller than the Chlorophyta dataset (303 genes vs. 2168 genes), which probably explains the differences in the fraction of genes found. Nevertheless, these results indicate that our *de novo* assembled transcriptome has captured the majority of the expressed genes in *Acetabularia*.

**Table 2.**
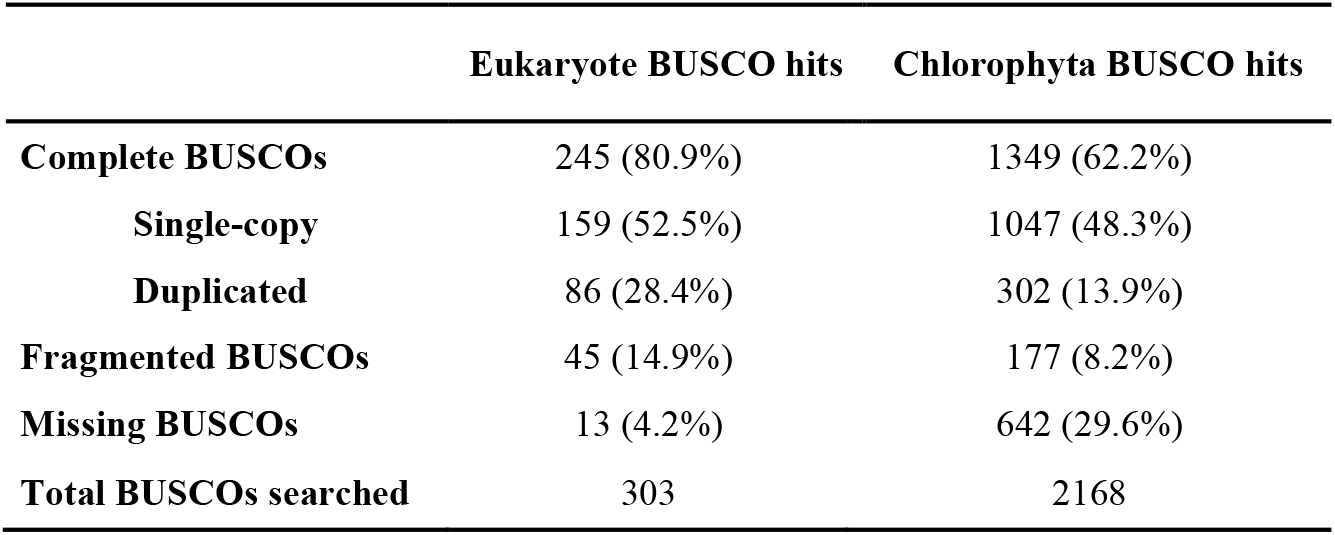
BUSCO analysis of the *de novo* assembled transcriptome of *A. acetabulum*. BUSCOs refer to the genes present in the different databases of the BUSCO software. Two datasets were used in our analysis, one containing 2168 genes conserved across Chlorophyta, and one containing 303 genes conserved across eukaryotes.

|  | Eukaryote BUSCO hits | Chlorophyta BUSCO hits |
| --- | --- | --- |
| **Complete BUSCOs** | 245 (80.9%) | 1349 (62.2%) |
| **Single-copy** | 159 (52.5%) | 1047 (48.3%) |
| **Duplicated** | 86 (28.4%) | 302 (13.9%) |
| **Fragmented BUSCOs** | 45 (14.9%) | 177 (8.2%) |
| **Missing BUSCOs** | 13 (4.2%) | 642 (29.6%) |
| **Total BUSCOs searched** | 303 | 2168 |

### RNA distribution

For all 24 samples, more than 70% of the trimmed read pairs mapped to the transcriptome (Table 1). There was no correlation between the number of mapped reads and the total UMI count (Table 1), illustrating the extent of PCR duplication in the sequence libraries and the importance of using UMIs. The majority of expressed transcripts were lowly expressed and 40-50% of the expressed transcripts had a count of two or less.

The total UMI counts follow the same distribution as the amount of isolated total RNA, with highest numbers in the cap samples and decreasing towards the rhizoid (Figure 1B and C). As the dissected cap pieces were larger than the other pieces, it is also expected that the cap samples contain the most RNA. However, as the same amount of RNA is sequenced from each library, the size of the pieces, or the amount of isolated total RNA, cannot explain the higher count values in the cap. This rather indicates a greater diversity, or heterogeneity, of transcripts in the cap libraries compared to the other libraries.

While the nuclear encoded mRNAs had the same distribution as the total RNA, i.e. decreasing towards the rhizoid (Figure S1A), ribosomal RNAs were roughly evenly distributed between the different samples, although with a few extreme outliers (Figure S1B). Transcripts presumably originating from the chloroplast were also distributed in an apical-basal gradient (Figure S1C). Mitochondrial transcripts had the highest counts in the cap and the rhizoid (Figure S1D), however these genes were much more variable between the samples and the counts were also overall much lower and therefore more subjected to stochastic variation.

The principal component analysis (PCA) of the count variation between samples shows that the samples largely cluster according to the cell apical-basal axis along PC1 (Figure 1D). This shows that the cap- and the rhizoid are the least similar in transcript composition, while the two stalk samples largely overlap and are fairly similar in terms of transcript expression. However, there also seem to be a slight tendency that the samples cluster along PC2 according to which batch they originate from (e.g. batch 19) or which RNA isolation method was used (e.g. batch 19 and 25 vs. batch 26, 27, 25 and 46). This indicates that also which batch of cells the samples originated from (i.e. sampled at the same time), or which RNA isolation kit was used also affects the sample variation.

### Differential transcript distribution

The majority of the assembled transcripts were expressed at low levels (which is expected as transcriptomes assembled *de novo* from NGS data always contains a high number of assembly artefacts and wrongly assembled isoforms). 87% of the transcripts had a mean expression of >1 TMM across the samples, and filtering nuclear encoded transcripts with a raw count of one or more in at least four samples retained 82,164 transcripts. Out of these, 13,057 transcripts were identified as significantly differentially distributed between at least two subcellular compartments. Of the differentially distributed transcripts, 4,197 transcripts were uniquely located in the rhizoid, 2,710 transcripts were unique to the cap, and 255 and 259 transcripts were uniquely located in the upper- and lower stalk respectively (Figure 2A). 1,617 transcripts were enriched in both the cap and the upper stalk, 163 transcripts in both the upper- and lower stalk, and 1,234 transcripts were enriched in both the lower stalk and the rhizoid. Visualizing the expression of these differentially distributed transcripts (Figure 2B) confirms the clustering analysis in that there are two large and distinct pools of enriched transcripts in the cap and the rhizoid, and that these subcellular compartments are the least similar in terms of gene content. The upper- and lower stalk samples have similar expression profiles and share a large number of differentially distributed transcripts. These two compartments also have an overall lower gene expression compared to the cap and rhizoid.

**Figure 2.**
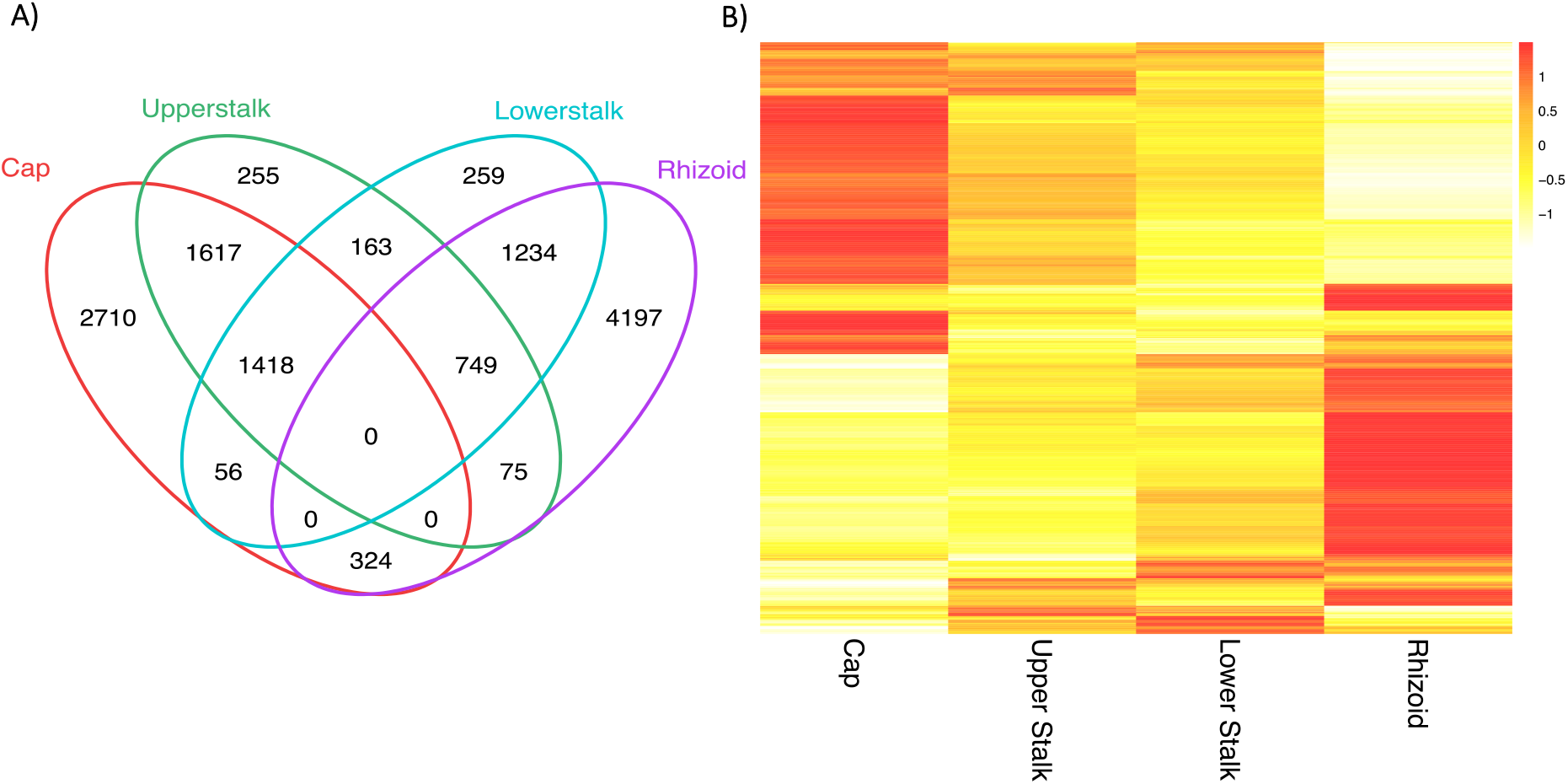
Differentially distributed transcripts between the subcellular compartments. A) Venn diagram showing the shared and unique number of transcripts that are differentially distributed between the subcellular compartments. B) Heatmap of the differentially distributed transcripts. Colors represent scaled TMM (Trimmed Mean of M-values) expression values. The mean TMM values across the different samples from each subcellular structure is shown.

### GO enrichment

In order to investigate which genetic processes were taking place in the different subcellular compartments, we analyzed the different subcellular pools of nuclear encoded transcripts for the presence of enriched functional categories. As the two stalk samples displayed very similar expression patterns, they were analyzed together (referred to as “stalk”) to get a clearer picture of the differences between the stalk, cap and the rhizoid. The GO-enrichment analysis resulted in 126 enriched GO-terms in the cap, 134 in the rhizoid, and 57 in the stalk (there were eight enriched GO-terms in the upper stalk and 14 in the lower stalk when analyzed separately) (Table S7).

Nuclear encoded mRNA transcripts accumulating in the cap were enriched for GO-terms related to photosynthesis such as photosynthetic processes, chloroplast components and thylakoid (Figure 3). General metabolic processes, organization of the plasma membrane and extracellular matrix, development and transport were also enriched in the cap, and to a lesser extent enriched in the rhizoid. No particular processes seemed to be unique to the stalk. However, the GO-term “cytoplasmic chromosome”, which was also enriched in the cap, was significantly enriched in the stalk, and GO-terms related to metabolic processes, catalytic activity, cellular organelles and transport was to small degree enriched in the stalk. Nuclear encoded mRNA transcripts accumulating in the rhizoid were enriched for GO-terms related to the nucleus, replication, transcription, and cell motility. In addition, cytoskeleton organization and cell-division and differentiation were enriched in the rhizoid and also to a lesser extent in the cap. Other processes which seemed to be more widely distributed and which were enriched in both the cap and the rhizoid were related to, transport, translation, ribosome organization, cell wall- and cell membrane organization, development and morphogenesis, and general metabolic and enzymatic processes.

**Figure 3.**
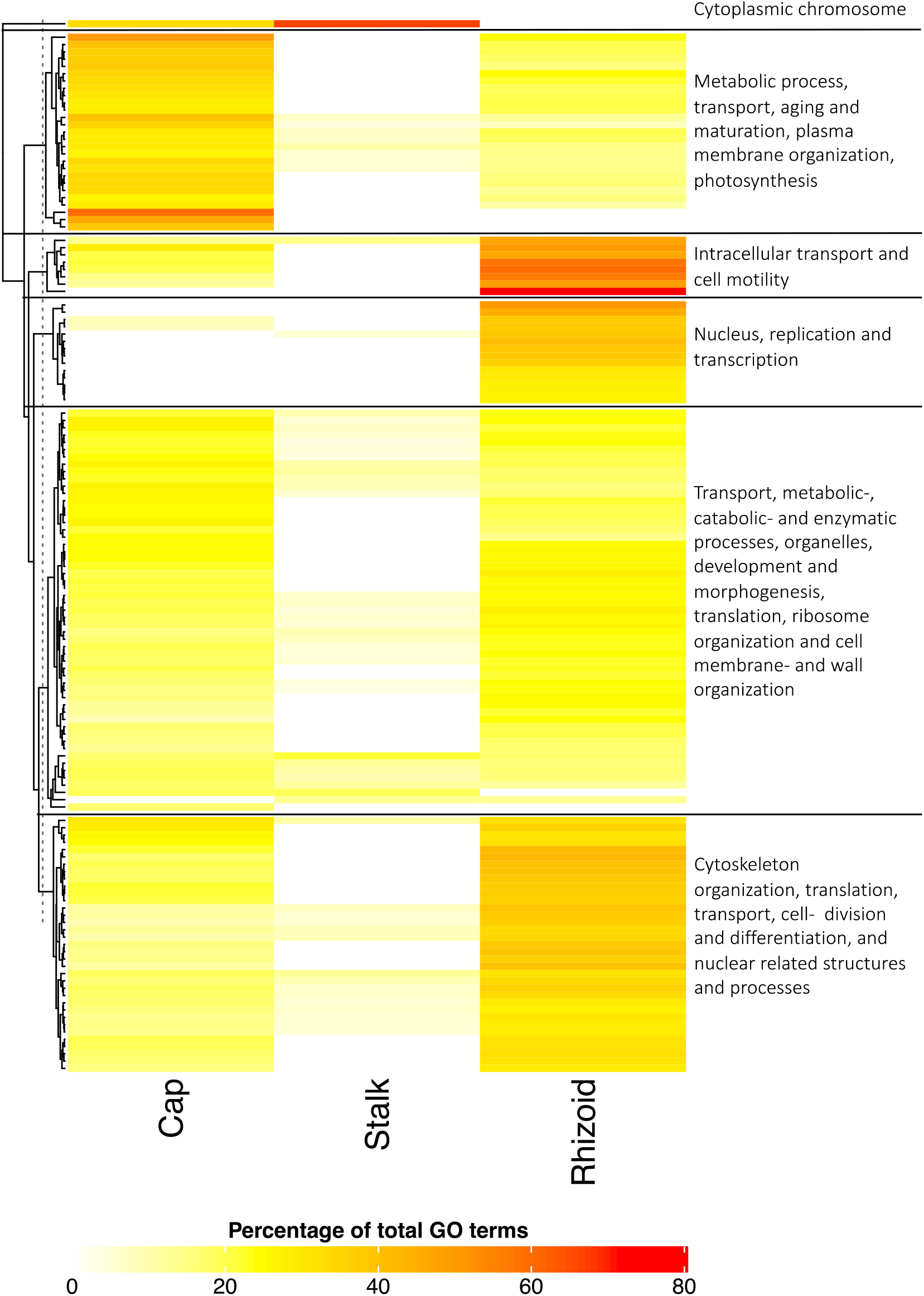
GO-enrichment analysis of transcripts differentially distributed between subcellular compartments. Heatmap of the enriched GO-terms among the differentially distributed transcripts in each subcellular compartment (note that the stalk samples are analyzed together). The colors indicate the percentage of differentially distributed transcripts annotated with a given GO-category compared to the total number of transcripts in the same GO-category. All GO-categories (Biological Process, Cellular Compartment and Molecular Function) are shown together. A GO-term not significantly enriched in a subcellular compartment is set to zero percent (hence shown in white color). The most prevalent GO-terms have been simplified and highlighted on the right side of the heatmap. See table S7 for a full description of the GO-enrichment results.

### Distribution of genes involved in mRNA compartmentalization

Analyses of the mRNA distribution indicated the presence of functionally related subcellular pools of transcripts. Therefore, we investigated in detail the distribution patterns of transcripts potentially involved in generating this type of distribution.

Transcripts related to the cytoskeleton, such as actin and tubulin, are highly abundant along the entire cell, and apparently not specifically associated with any particular subcellular region (Figure 4A and B). Transcripts encoding motor proteins moving along the cytoskeleton such as myosin, dynein and kinesin, are also present throughout the cell, although not as evenly distributed as the cytoskeletal components (Figure 4C-E). Myosin transcripts were more abundant in the apical end and decreasing towards the rhizoid. This includes class XIII myosin which have been shown to be involved in organelle transport and tip growth in *A. cliftonii* and enriched in the apical regions of the cell (Vugrek et al., 2003). The same trend was observed also for kinesins, except for two transcripts which were most abundant in the rhizoid. Interestingly, these two transcripts have the closest blast hits against kinesin 13 and 14, which are known to move in both directions on the microtubule, and can thereby travel in the opposite direction on the microtubules than the other kinesins.

**Figure 4.**
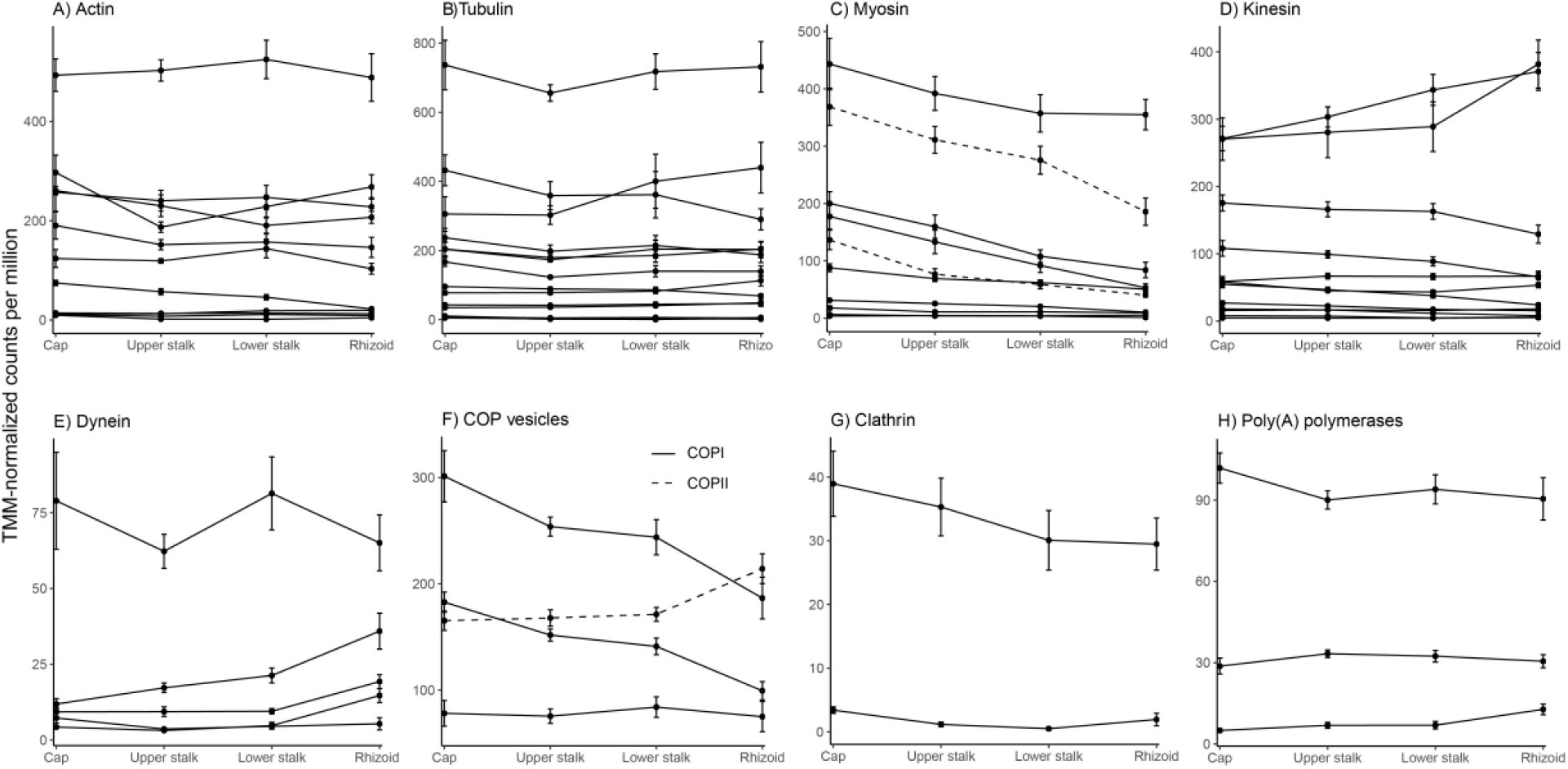
Distribution of transcripts related to transport and RNA localization. The distribution of homologs of Actin (A), Tubulin (B), Myosin (C) – the dashed lines indicates homologs of Class XIII myosin identified in *A. cliftonii*, Kinesin (D), Dynein (E), genes creating COP vesicles (F), Clathrin (G) and poly(A) polymerases (H). Expression values (TMM-normalized counts per million) are shown on the y-axis, with error bars representing standard error. Only transcripts with a mean normalized expression >1 across all samples are shown.

In contrast, the dyneins were overall lesser expressed than myosins and kinesins. The most highly abundant dynein was present through the cell in roughly equal amounts, while the rest of the transcripts were most abundant in the rhizoid. Myosins and kinesis generally move towards the plus-ends of the polarized actin microfilaments and microtubules respectively, and thus from the nucleus towards the cell membrane. While dyneins move toward the minus end of microtubules towards the cell interior (Alberts et al., 2002). Hence, motor proteins moving towards the cell membrane are of a slightly higher abundance in the apical part of the cell (expect from two kinesin transcripts which are of a higher abundance in the basal part of the cell), while motor proteins moving towards the cellular interior are seemingly of a higher abundance in the basal part of the cell (lower stalk and rhizoid).

Vesicular transport is a fundamental mechanism for intracellular transport of cargo in eukaryote cells, and has been associated with intracellular transport of RNA (Basyuk et al., 2003; Roberts et al., 2013; Skog et al., 2008). Vesicle formation relies on coat proteins, and COPI-COPII- and Clathrin coated vesicles are the main type of vesicles in eukaryote cells. COPI-coated vesicles move from ER to golgi, COPII-coated vesicles move between parts of golgi and retrograde transport from golgi to ER, and Clathrin-coated vesicles move from golgi to the plasma membrane (Gomez-Navarro et al., 2016). In our results, two of three COPI transcripts were most abundant in the cap and decrease towards the rhizoid, while one COPII transcript was most abundant in the rhizoid (Figure 4F). Two Clathrin homologs were distributed throughout the cell, one of these transcripts was of noticeable higher abundance than the other (Figure 4G). This high abundant transcript was also of a slightly higher concentration in the cap and decreasing towards the rhizoid.

Three copies of poly(A) polymerases were expressed in *A. acetabulum*. All three were evenly distributed throughout the cell, however one was higher expressed than the others (Figure 4H).

### Comparative transcriptomics between *A. acetabulum* and *Caulerpa taxifolia*

Orthology searches between the transcriptomes of *A. acetabulum* and *Caulerpa taxifolia* resulted in 4,483 orthogroups represented by at least one transcript from each species. Of these orthogroups, 2,120 were single-copy orthologues with a single representative from each species. Comparison of the different samples from the two species based on expression dynamics of these single-copy orthologues showed that the genes clustered strongly according to species, rather than to subcellular compartment between species (Figure 5).

**Figure 5.**
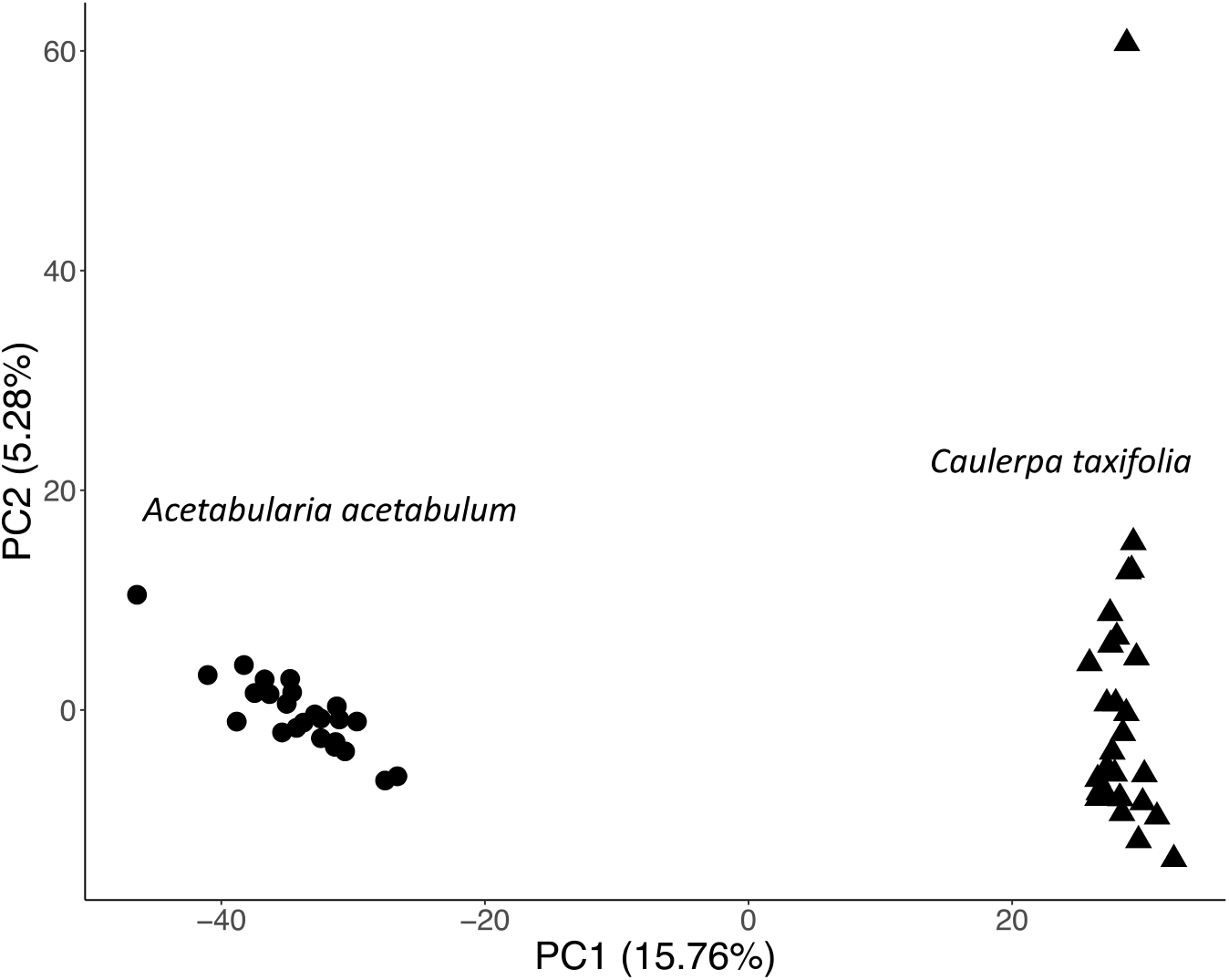
Comparison of the expression profile of gene orthologues between *A. acetabulum* and *C. taxifolia*. Principal Component Analysis (PCA) of the sample variation based on TMM-normalized counts per million (see Methods) of single-copy orthologues between *A. acetabulum* (circles) and *C. taxifolia* (triangles).

## Discussion

### Apical-basal mRNA gradient in *A. acetabulum*

*Acetabularia acetabulum* has been used as a model system for cell morphogenesis for decades, and it has been suspected that differential distribution of RNA long the cell axis is an underlying mechanism for its sophisticated morphology (Dumais et al., 2000; Hämmerling, 1934b; Serikawa et al., 2001; Vogel et al., 2002). To investigate the distribution of mRNA in adult *A. acetabulum* cells we have performed subcellular mRNA sequencing and functional enrichment analysis. We have tagged each mRNA molecule with unique molecular indexes (UMIs) which allows for true quantification of mRNA by eliminating the effect of amplification bias introduced during library preparation. Despite isolating RNA from subsections of a single-cell, we were able to capture the majority of expressed transcripts, and although there is some variation between samples the procedure was repeatable and robust even to the type of RNA isolation protocols.

Our results demonstrate the presence of RNA throughout the entire cell length, and identified the highest amount of mRNA at the apical end of the cell (the cap) decreasing towards the basal end (the rhizoid), confirming earlier discoveries of an apical-basal gradient of RNA in *A. acetabulum* (Baltus et al., 1968; Hämmerling, 1936; Werz, 1955). However, while it has been believed that this gradient is due to different concentrations of RNA encoded by the chloroplasts, and not nuclear encoded RNAs (Naumova et al., 1976), we rather find that the apical-basal gradient is mainly caused by nuclear encoded mRNAs.

### Localized pools of transcripts support subcellular mRNA compartmentalization

A long-standing question has been whether the observed gradient of mRNA in *A. acetabulum* is homogeneous in transcript composition, or whether there are distinct pools of transcripts along the cell. Hämmerling’s grafting experiments suggested the existence of local determinants of morphogenesis, and Dumais et al. (2000) speculated that mRNAs would either be distributed throughout the cell, or localized to the apical or basal ends. Our results show that while some gene transcripts are distributed evenly across the entire cell, a large part are actually abundantly located to different subcellular compartments. We also found that these pools of transcripts are composed of functionally related transcripts. Transcripts related to photosynthesis are co-localized and accumulated in the apical end of the cell, while transcripts related to nuclear processes co-localized in the basal end. This pattern shows that the RNA gradient is not a homogeneous mix of gene transcripts, which confirms that mechanisms to ensure specific and functional RNA localization must be in place in *A. acetabulum*.

There were overall fewer transcripts localized in the stalk. This was not surprising as the stalk is mainly filled with a central vacuole, with only a thin layer of cytoplasm covering it (Dumais et al., 2000), leaving very little room for other subcellular structures or pools of transcripts. We therefore assume that there are very few processes happening exclusively in the stalk and that the stalk might even function as a physical border between the cap and the rhizoid, making sure that molecules and other substances are not mixed between the two compartments, but instead carefully transported between them.

### Active mRNA transport is likely the main mechanism for establishment of cell polarity

RNA can be distributed around a cell either by passive diffusion from the nucleus, or by active transport along the cytoskeleton (St Johnston, 2005). Studies tracking the movement of radioactive labelled RNA labelled have shown that mRNA travel faster in *A. acetabulum* than what is possible by diffusion alone (Kloppstech et al., 1975b), and simply the size of the cell, with the nucleus and the cap separated by several centimeters, puts obvious demands on active intracellular transport. A highly sophisticated and extensively developed cytoskeleton has been overserved in *A. acetabulum*, with large tracks of actin filaments running the entire length of the cell (Menzel, 1994). Experiments by Mine et al. (2001) showed that inhibiting actin polymerization with cyclohalasin D disrupts the established mRNA gradients, indicating that there is an association between mRNA and the cytoskeleton. As expected, we find that transcripts encoding the main cytoskeletal components such as actin and tubulin are uniformly distributed throughout the cell. Furthermore, we see that both Clathrin and COP genes, as well as homologs of motor proteins traveling in both directions on the cytoskeleton, are distributed throughout the cell, suggesting that these types of vesicular transport systems are active in the entire cell.

### mRNA stabilization and post-transcriptional control

The fact that some transcripts are evenly distributed while others are localized, implies that the cell is either able to distinguish between which mRNAs should be transported where, but can also mean that there are mechanisms for selective stabilization and degradation of mRNAs at different locations in the cell. While actin microfilaments are present throughout the cell for the entire life cycle of *A. acetabulum*, microtubules do not appear until the final stages of development where they serve as transport tracks in the cap (Menzel, 1986; Menzel, 1994). This is interesting as tubulin genes are expressed much earlier and distributed throughout the cell, so tubulin mRNAs must be stabilized and stored in the cytoplasm and prevented from being translated. That mRNA is long-lived in *A. acetabulum* cells is supported by experiments showing that development and morphogenesis can continue for days, and even weeks, after amputation of the nucleus (Stich et al., 1958; Yasinovski et al., 1979), and that radioactive RNAs exist in *A. acetabulum* cells long after treatment with radioactive labelled UTP (Kloppstech et al., 1982; Schweiger, 1977). It is also interesting that we find polyadenylation genes distributed throughout the cell, as editing, shortening and elongation of the poly-A-tail is an important mechanism for translational control (Aphasizhev, 2005; Wickens, 1992).

### The cap is the main morphogenetic and metabolic structure

The GO-enrichment analysis also shows a higher level of catalytic- and metabolic activity in the cap compared the rhizoid, indicating the cap is the metabolically more diverse and active region of the cell. We also see the greatest diversity of expressed transcripts in the cap, and the highest overall RNA content. These finding agree with earlier observations that the cap is the section of the cell with the highest morphogenetic capacity, or developmental potential, as it can regenerate both whorls of hair and the entire cap structure after being dissected from the rhizoid (Menzel, 1994; Serikawa et al., 2001), while the nucleus is mostly the place for production of mRNAs and replication.

### Ortholog comparison indicates little genetic homology between subcellular sections of *A. acetabulum* and *Caulerpa taxifolia*

*Caulerpa taxifolia* is another Chlorophyte algae with many similarities to *A. acetabulum*, most notably they are both gigantic single-celled species with highly complex cellular morphologies with clearly distinguished apical and basal ends. However, unlike *A. acetabulum*, *C. taxifolia* is a syncytium with hundreds, or even thousands, of nuclei scattered throughout the cell. Ranjan et al. (2015) characterized the gene expression patterns of the different subcellular sections of *C. taxifolia* and found that they contained unique expression profiles, similar to what we see in *A. acetabulum*. However, comparing the expression profile of single-copy orthologs between the two species shows that the subcellular sections are more similar within each species rather than between species, indicating that there are different genes active in the apical and basal sections in these two species, and hence little homology at the genetic level. Nevertheless, comparing the functional annotations of these genes indeed shows some similarities. Both species are enriched for nuclear components and DNA-related processes, such as DNA replication and transcription in the basal parts of the algae, as well as displaying a higher catalytic activity in the apical parts (Ranjan et al., 2015). Therefore, although morphologically similar cell sections of these two species contain largely non-orthologous mRNAs, they do seem to share overall similar genetic functions. This might be an indication of evolutionary convergence on a functional level in the two species.

## Supplementary information

### Supplementary tables

**Table S1.**
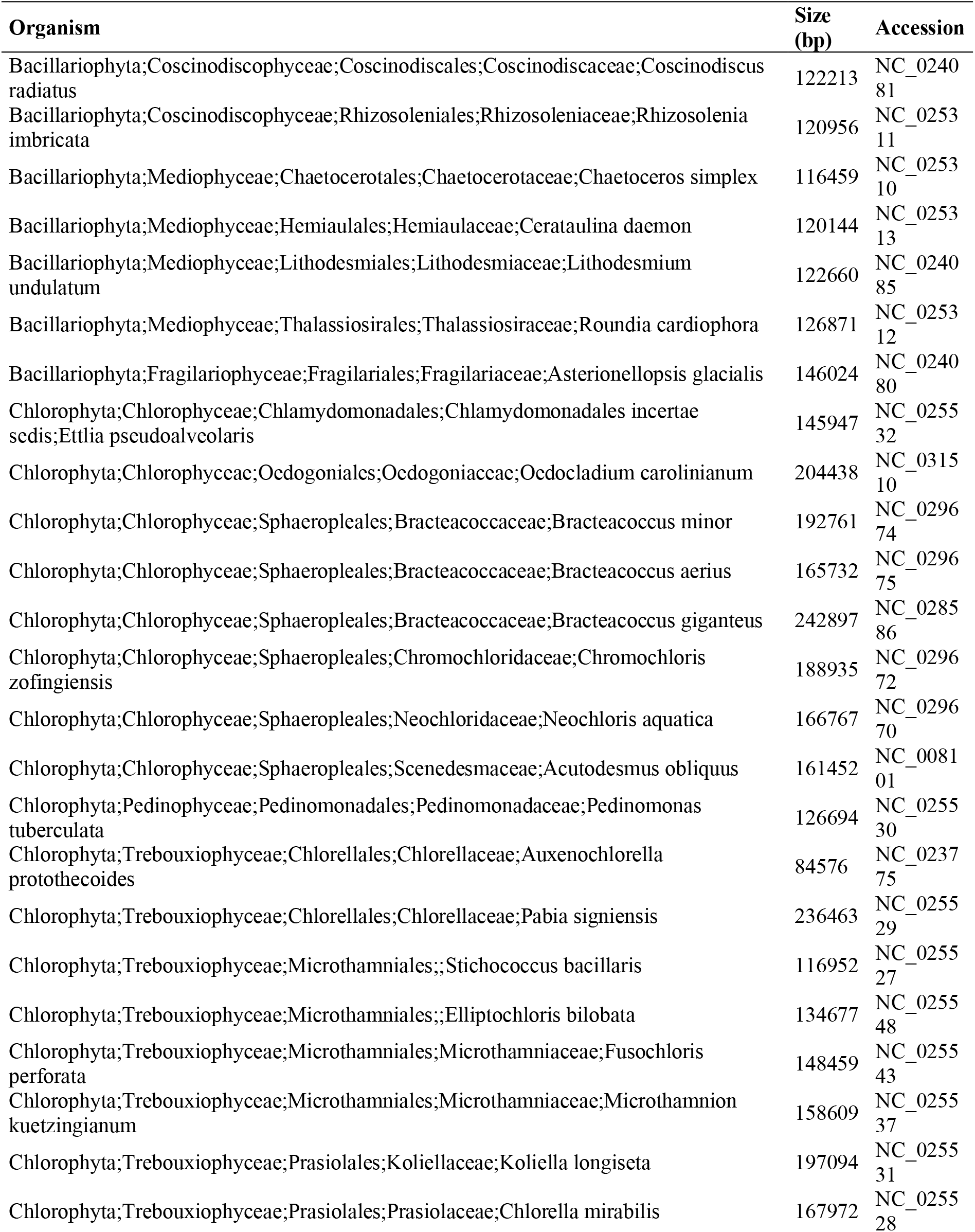
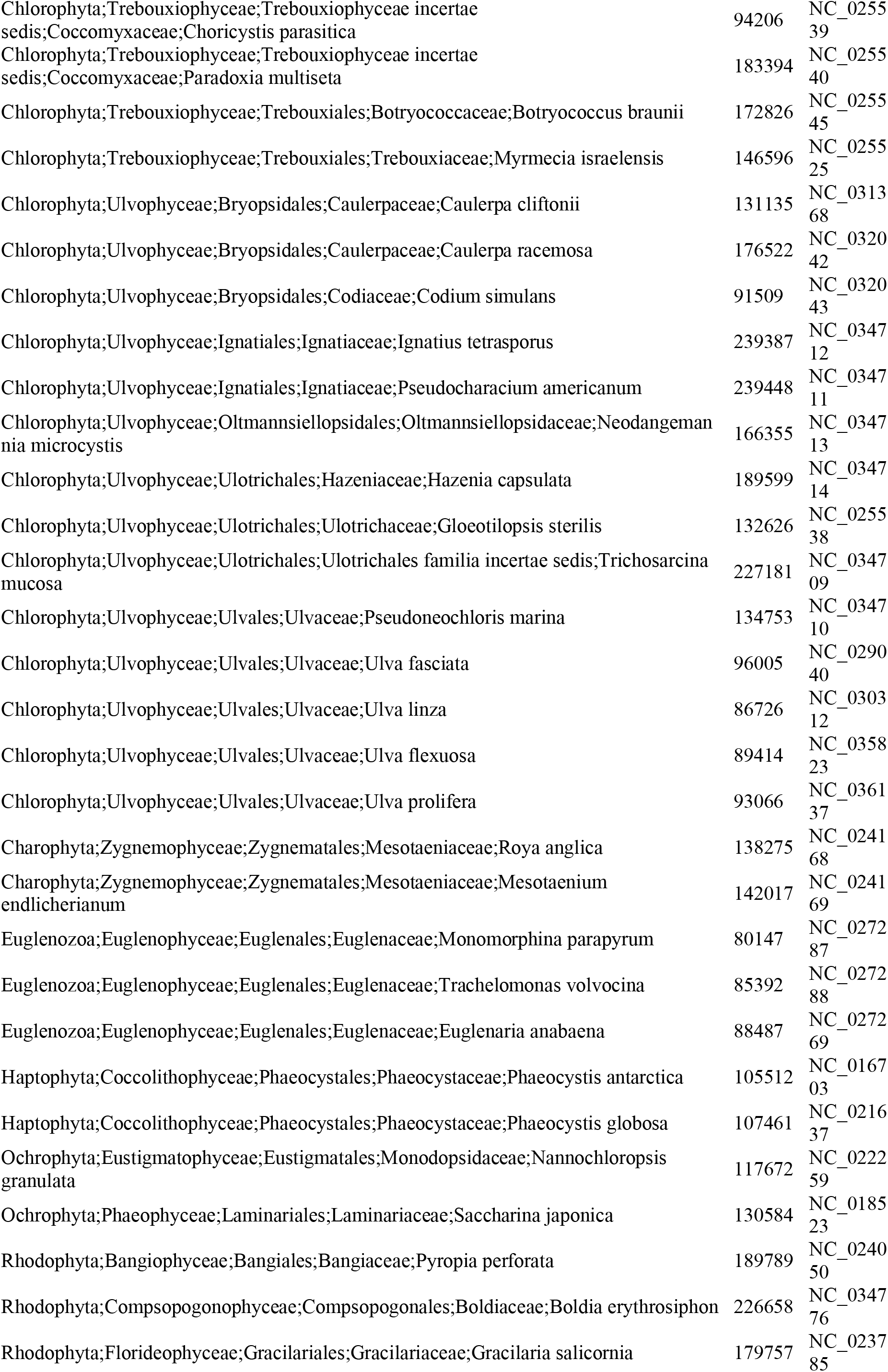
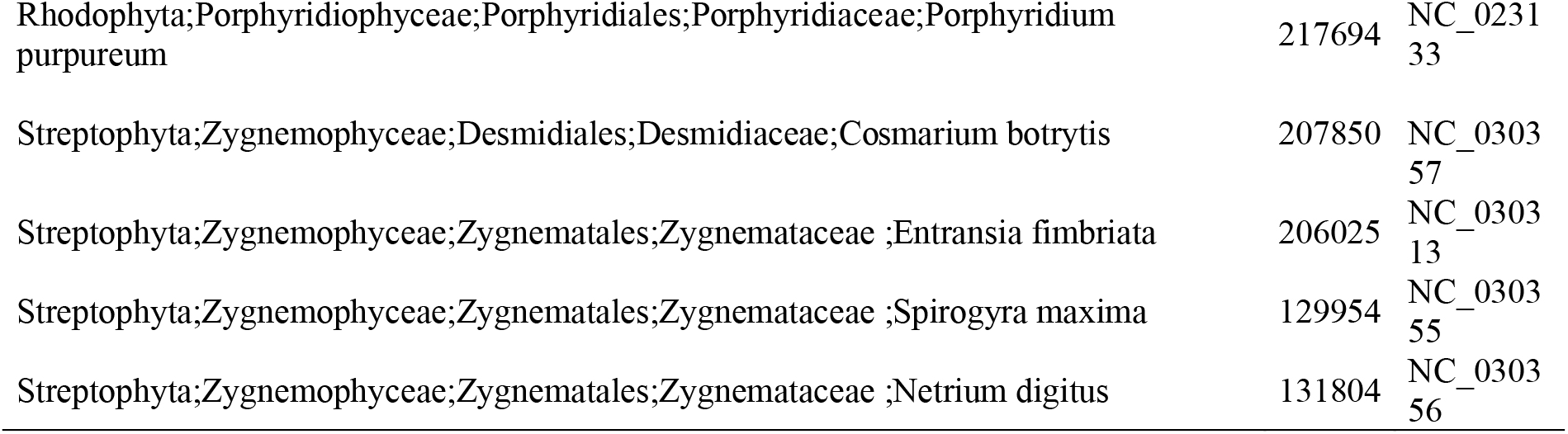
Chloroplast genomes used as BLAST database to identify *Acetabularia acetabulum* chloroplast transcripts.

**Table S2.** Mitochondrial genomes used as BLAST database to identify *Acetabularia acetabulum* mitochondrial transcripts

| Organism | Size (bp) | Accession |
| --- | --- | --- |
| Eukaryota;Stramenopiles;PX clade;Phaeophyceae;Laminariales;Laminariaceae;Saccharina;Saccharina latissima | 3765<br>9 | NC_02<br>6108 |
| Eukaryota;Sar;Stramenopiles;Ochrophyta;PX clade;Phaeophyceae;Laminariales;Laminariaceae;Saccharina;Saccharina japonica | 3765<br>7 | NC_01<br>3476 |
| Eukaryota;Stramenopiles;Eustigmatophyceae;Eustigmatales;Monodopsidaceae;Nannochloropsis;Nannochloropsis oceanica | 3805<br>7 | NC_02<br>2258 |
| Eukaryota;Rhodophyta;Florideophyceae;Rhodymeniophycidae;Gracilariales;Gracilariaceae;Gracilaria;Gracilaria salicornia | 2527<br>2 | NC_02<br>3784 |
| Eukaryota;Viridiplantae;Streptophyta;Zygnemophyceae;Zygnematales;Mesotaeniaceae;Roya;Roya obtusa | 6946<br>5 | NC_02<br>2863 |
| Eukaryota;Rhodophyta;Bangiophyceae;Bangiales;Bangiaceae;Pyropia;Pyropia nitida | 3531<br>3 | NC_02<br>7616 |
| Eukaryota;Viridiplantae;Chlorophyta;Trebouxiophyceae;Trebouxiophyceae incertae sedis;Botryococcaceae;Botryococcus;Botryococcus braunii | 8458<br>3 | NC_02<br>7722 |
| Eukaryota;Viridiplantae;Chlorophyta;Trebouxiophyceae;Chlorellales;Chlorellaceae;Chlorella | 5252<br>8 | NC_02<br>4626 |
| Eukaryota;Viridiplantae;Chlorophyta;Trebouxiophyceae;Chlorellales;Chlorellaceae;Auxenochlorella;Auxenochlorella protothecoides | 5727<br>4 | NC_02<br>6009 |
| Eukaryota;Viridiplantae;Chlorophyta;core chlorophytes;Trebouxiophyceae;Chlorellales;Chlorellaceae;Chlorella clade;Chlorella;Chlorella variabilis | 7850<br>0 | NC_02<br>5413 |
| Eukaryota;Viridiplantae;Chlorophyta;Pedinophyceae;Pedinomonadales;Pedinomonadaceae;Pedinomonas;Pedinomonas minor | 2513<br>7 | NC_00<br>0892 |
| Eukaryota;Viridiplantae;Chlorophyta;Chlorophyceae;Sphaeropleales;Neochloridaceae;Neochloris;Neochloris aquatica | 3802<br>1 | NC_02<br>4761 |
| Eukaryota;Viridiplantae;Chlorophyta;Chlorophyceae;Sphaeropleales;Chromochloridaceae;Chromochloris;Chromochloris zofingiensis | 4484<br>0 | NC_02<br>4758 |
| Eukaryota;Viridiplantae;Chlorophyta;Chlorophyceae;Sphaeropleales;Neochloridaceae;Chlorotetraedron;Chlorotetraedron incus | 3840<br>6 | NC_02<br>4757 |
| Eukaryota;Viridiplantae;Chlorophyta;Chlorophyceae;Sphaeropleales;Bracteacoccaceae;Bracteacoccus;Bracteacoccus aerius | 4715<br>8 | NC_02<br>4755 |
| Eukaryota;Viridiplantae;Chlorophyta;Chlorophyceae;Sphaeropleales;Bracteacoccaceae;Bracteacoccus;Bracteacoccus minor | 4517<br>5 | NC_02<br>4756 |
| Eukaryota;Viridiplantae;Streptophyta;Zygnemophyceae;Zygnematales;Zygnemataceae;Entransia;Entransia fimbriata | 6164<br>5 | NC_02<br>2861 |
| Eukaryota;Viridiplantae;Chlorophyta;Ulvophyceae;OUU clade;Ulvaes;Ulvaceae;Ulva;Ulva fasciata | 6161<br>4 | NC_02<br>8081 |
| Eukaryota;Viridiplantae;Chlorophyta;Ulvophyceae;OUU clade;Ulvaes;Ulvaceae;Ulva;Ulva prolifera | 6384<br>5 | NC_02<br>8538 |
| Eukaryota;Viridiplantae;Chlorophyta;Ulvophyceae;OUU clade;Ulvaes;Ulvaceae;Ulva;Ulva linza | 7085<br>8 | NC_02<br>9701 |
| Eukaryota;Viridiplantae;Chlorophyta;Ulvophyceae;OUU clade;Ulvaes;Ulvaceae;Ulva;Ulva pertusa | 6933<br>3 | NC_03<br>5722 |
| Eukaryota;Viridiplantae;Chlorophyta;Ulvophyceae;OUU clade;Ulvaes;Ulvaceae;Ulva;Ulva flexuosa | 7154<br>5 | NC_03<br>5809 |
| Eukaryota;Viridiplantae;Chlorophyta;Chlorophyceae;Sphaeropleales;Selenastraceae;Ourococcus;Ourococcus multispurus | 4970<br>5 | NC_02<br>4762 |
| Eukaryota;Viridiplantae;Chlorophyta;Chlorophyceae;Chlamydomonadales;Chlamydomonada ceae;Polytoma;Polytoma uvella | 1741<br>1 | NC_02<br>6572 |

**Table S3.**
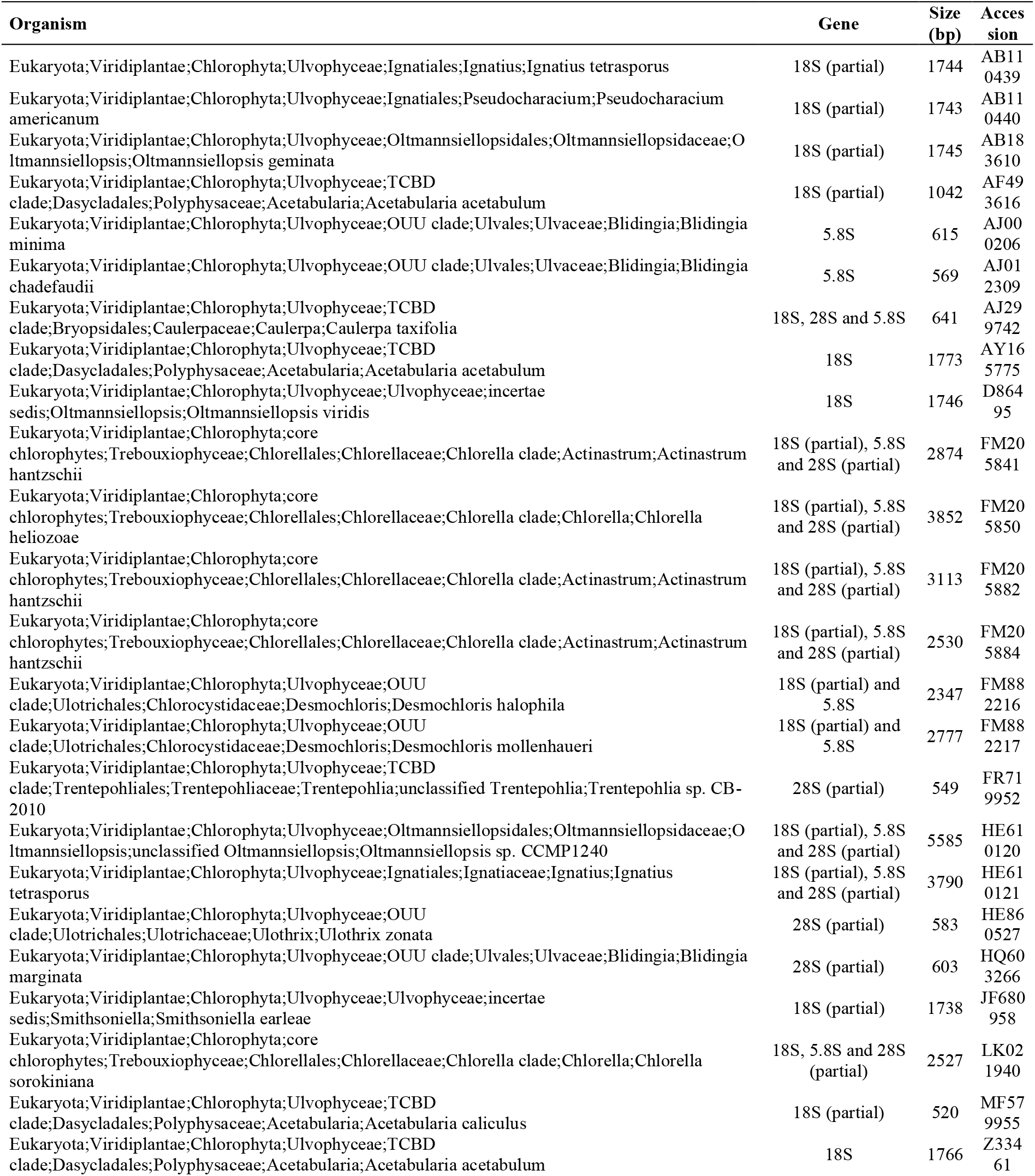
rRNA sequences from various green algae used as BLAST database to identify *Acetabularia* rRNA transcripts.

**Table S4.** *Chlamydomonas reinhardthii* COP and Clathrin genes used to identify similar genes in *A. acetabulum*

| Gene | Accession number |
| --- | --- |
| alpha-COP | XP_001693686 |
| beta-COP | XP_001701255 |
| beta-COP | XP_001702422 |
| delta-COP | XP_001701763 |
| epsilon-COP | XP_001693085 |
| gamma-COP | XP_001691193 |
| zeta-COP, subunit of COP-1 complex | XP_001701294 |
| sar-type small GTPase | XP_001699537 |
| COP-II coat subunit | XP_001700438 |
| COP-II coat subunit | XP_001702936 |
| COP-II coat subunit | XP_001701974 |
| COP-II coat subunit | XP_001697039 |
| Clathrin heavy chain | XP_001699806 |

**Table S5.** RNA isolation.

|  | Single Cell RNA<br>purification Kit | Total RNA<br>purification kit | % Increase by using the<br>Total RNA purification<br>kit |
| --- | --- | --- | --- |
| Mean total RNA yield (ng) | 33 | 159 | 382 |
| Mean transcripts with count > 0 | 51715 | 55031 | 6 |
| Mean transcripts with count > 1 | 35019 | 37170 | 6 |
| Mean transcripts with count > 2 | 28821 | 30510 | 6 |
| Mean transcripts with count > 3 | 25474 | 26897 | 6 |

**Table S6.** Assembly statistics of the *de novo* assembled transcriptome of *Acetabularia acetabulum*. Only the highest expressed isoform of each gene is represented in the transcriptome. This transcriptome was used as a mapping reference for further RNA-seq analysis.

| *A. acetabulum* transcriptome |  |
| --- | --- |
| Total transcriptome size (bp) | 114 907 754 |
| Number of transcripts | 246 083 |
| Longest transcript (bp) | 17 196 |
| Shortest transcript (bp) | 201 |
| Mean transcript length (bp) | 467 |
| Median transcript length (bp) | 298 |
| N50 transcript length (bp) | 534 |
| Nuclear encoded transcripts | 245 334 |
| mRNAs | 245 223 |
| rRNAs | 111 |
| Chloroplast encoded transcripts | 389 |
| Mitochondrial encoded transcripts | 99 |
| Possible prokaryotic/unclassified transcripts | 261 |
| Predicted protein coding transcripts (TransDecoder) | 114 146 |
| Nuclear | 113 900 |
| Chloroplast | 178 |
| Mitochondrial | 68 |

**Table S7.** Enriched Gene Ontologies. Domain refers to “Cellular Component” (CC), “Biological Process” (BP) and “Molecular Function” (MF). FDR refers to the False Discovery Rates calculated on the p-values by the qvalue function in the R package qvalue. The “%” in the Cap, Stalk and Rhizoid is the percentage of GO-ID’s in the enriched pool of transcripts (in the Cap, Stalk or Rhizoid) compared to the number in the total transcriptome.

| ID | Term | Domain | Cap<br>FDR | Cap<br>% | Stalk<br>FDR | Stalk<br>% | Rhizoid<br>FDR | Rhizoid<br>% |
| --- | --- | --- | --- | --- | --- | --- | --- | --- |
| GO:0005623 | cell | CC | 4.3E-181 | 20 | 4.9E-34 | 5 | 1.1E-208 | 23.2 |
| GO:0009536 | plastid | CC | 2.2E-103 | 35 | 3.3E-04 | 4 | 1.5E-02 | 8.1 |
| GO:0009579 | thylakoid | CC | 1.8E-60 | 47 | 0 | 0 | 0 | 0 |
| GO:0006950 | response to stress | BP | 1.0E-40 | 23 | 3.8E-06 | 5 | 8.3E-30 | 22 |
| GO:0009058 | biosynthetic process | BP | 1.5E-37 | 23 | 4.6E-07 | 6 | 1.3E-32 | 23 |
| GO:0005829 | cytosol | CC | 2.7E-31 | 19 | 6.0E-06 | 4 | 1.3E-47 | 24 |
| GO:0043167 | ion binding | MF | 5.3E-28 | 26 | 0 | 0 | 6.4E-17 | 22 |
| GO:0005886 | plasma membrane | CC | 2.5E-25 | 22 | 6.3E-03 | 4 | 1.1E-24 | 24 |
| GO:0022857 | transmembrane transporter activity | MF | 2.5E-25 | 40 | 4.0E-03 | 7 | 9.9E-03 | 12 |
| GO:0044281 | small molecule metabolic process | BP | 5.4E-25 | 29 | 2.8E-05 | 7 | 5.0E-10 | 20 |
| GO:0016491 | oxidoreductase activity | MF | 5.7E-23 | 34 | 0 | 0 | 2.3E-03 | 13 |
| GO:0005773 | vacuole | CC | 3.7E-21 | 27 | 2.4E-03 | 5 | 1.3E-05 | 15 |
| GO:0051186 | cofactor metabolic process | BP | 2.0E-20 | 34 | 1.1E-02 | 6 | 7.7E-04 | 14 |
| GO:0015979 | photosynthesis | BP | 8.1E-20 | 38 | 0 | 0 | 0 | 0 |
| GO:0005739 | mitochondrion | CC | 9.3E-20 | 24 | 2.1E-02 | 4 | 4.0E-11 | 20 |
| GO:0032991 | protein-containing complex | CC | 8.8E-18 | 14 | 3.7E-07 | 5 | 3.9E-86 | 29 |
| GO:0065003 | protein-containing complex assembly | BP | 2.3E-17 | 26 | 0 | 0 | 2.1E-22 | 31 |
| GO:0006091 | generation of precursor metabolites and energy | BP | 3.8E-17 | 29 | 0 | 0 | 3.2E-03 | 11 |
| GO:0042592 | homeostatic process | BP | 1.9E-16 | 28 | 0 | 0 | 2.2E-08 | 21 |
| GO:0006520 | cellular amino acid metabolic process | BP | 6.0E-16 | 36 | 0 | 0 | 3.4E-05 | 20 |
| GO:0005576 | extracellular region | CC | 1.0E-15 | 31 | 2.0E-02 | 6 | 1.0E-03 | 14 |
| GO:0006629 | lipid metabolic process | BP | 1.2E-15 | 29 | 6.7E-03 | 6 | 1.0E-05 | 19 |
| GO:0005794 | Golgi apparatus | CC | 1.4E-15 | 26 | 5.2E-03 | 6 | 1.6E-05 | 17 |
| GO:0016887 | ATPase activity | MF | 4.2E-15 | 34 | 0 | 0 | 1.8E-03 | 16 |
| GO:0019899 | enzyme binding | MF | 6.7E-15 | 27 | 8.4E-04 | 7 | 1.8E-08 | 21 |
| GO:0005783 | endoplasmic reticulum | CC | 2.8E-13 | 24 | 0 | 0 | 1.3E-13 | 26 |
| GO:0005840 | ribosome | CC | 2.5E-12 | 15 | 0 | 0 | 4.2E-18 | 21 |
| GO:0006790 | sulfur compound metabolic process | BP | 3.0E-12 | 38 | 0 | 0 | 4.3E-03 | 16 |
| GO:0009056 | catabolic process | BP | 4.0E-12 | 23 | 3.8E-06 | 9 | 1.1E-06 | 18 |
| GO:0007165 | signal transduction | BP | 1.1E-11 | 18 | 7.7E-05 | 6 | 9.1E-21 | 26 |
| GO:0003735 | structural constituent of ribosome | MF | 3.7E-11 | 15 | 0 | 0 | 7.5E-14 | 18 |
| GO:0000003 | reproduction | BP | 3.9E-11 | 18 | 9.6E-04 | 5 | 7.4E-22 | 26 |
| GO:0034641 | cellular nitrogen compound metabolic process | BP | 4.6E-11 | 17 | 3.4E-03 | 4 | 1.7E-22 | 24 |
| GO:0061024 | membrane organization | BP | 1.6E-10 | 27 | 4.7E-02 | 5 | 2.7E-08 | 24 |
| GO:0006605 | protein targeting | BP | 1.6E-10 | 24 | 0 | 0 | 5.4E-07 | 20 |
| GO:0016192 | vesicle-mediated transport | BP | 1.8E-10 | 22 | 4.0E-06 | 9 | 3.6E-10 | 23 |
| GO:0048856 | anatomical structure development | BP | 1.9E-10 | 17 | 4.7E-08 | 8 | 5.0E-18 | 23 |
| GO:0006412 | translation | BP | 1.1E-09 | 15 | 2.2E-02 | 3 | 2.1E-20 | 24 |
| GO:0007568 | aging | BP | 1.6E-09 | 33 | 0 | 0 | 8.3E-05 | 22 |
| GO:0005618 | cell wall | CC | 8.4E-09 | 22 | 0 | 0 | 2.6E-05 | 17 |
| GO:0048646 | anatomical structure formation involved in morphogenesis | BP | 9.4E-09 | 24 | 0 | 0 | 9.0E-06 | 20 |
| GO:0009790 | embryo development | BP | 1.3E-08 | 23 | 0 | 0 | 2.3E-09 | 25 |
| GO:0002376 | immune system process | BP | 2.1E-08 | 20 | 3.2E-02 | 5 | 2.3E-13 | 28 |
| GO:0016829 | lyase activity | MF | 4.5E-08 | 32 | 0 | 0 | 9.3E-04 | 19 |
| GO:0006810 | transport | BP | 4.5E-08 | 35 | 0 | 0 | 9.6E-05 | 25 |
| GO:0006464 | cellular protein modification process | BP | 6.4E-08 | 15 | 9.6E-04 | 5 | 4.0E-31 | 29 |
| GO:0030154 | cell differentiation | BP | 8.7E-08 | 17 | 5.4E-03 | 5 | 1.0E-20 | 29 |
| GO:0008219 | cell death | BP | 1.4E-07 | 21 | 8.4E-03 | 7 | 4.0E-09 | 25 |
| GO:0003013 | circulatory system process | BP | 1.7E-07 | 60 | 0 | 0 | 0 | 0 |
| GO:0034655 | nucleobase-containing compound catabolic process | BP | 3.4E-07 | 18 | 1.2E-02 | 5 | 1.5E-09 | 23 |
| GO:0007267 | cell-cell signaling | BP | 4.4E-07 | 30 | 5.2E-03 | 11 | 2.3E-07 | 32 |
| GO:0005764 | lysosome | CC | 4.7E-07 | 29 | 5.2E-03 | 10 | 8.8E-03 | 15 |
| GO:0005975 | carbohydrate metabolic process | BP | 4.9E-07 | 20 | 3.8E-06 | 11 | 1.5E-04 | 17 |
| GO:0031410 | cytoplasmic vesicle | CC | 6.2E-07 | 27 | 1.2E-03 | 11 | 3.1E-04 | 20 |
| GO:0005768 | endosome | CC | 1.4E-06 | 20 | 7.1E-03 | 7 | 1.9E-09 | 25 |
| GO:0005634 | nucleus | CC | 1.8E-06 | 14 | 8.7E-04 | 5 | 3.3E-33 | 30 |
| GO:0055085 | transmembrane transport | BP | 2.6E-06 | 26 | 0 | 0 | 1.1E-03 | 19 |
| GO:0016301 | kinase activity | MF | 3.4E-06 | 20 | 0 | 0 | 4.5E-07 | 23 |
| GO:0015031 | protein transport | BP | 5.8E-06 | 18 | 4.0E-03 | 7 | 9.1E-17 | 33 |
| GO:0007010 | cytoskeleton organization | BP | 1.0E-05 | 18 | 0 | 0 | 4.6E-28 | 43 |
| GO:0005856 | cytoskeleton | CC | 1.1E-05 | 18 | 0 | 0 | 1.3E-20 | 39 |
| GO:0021700 | developmental maturation | BP | 1.4E-05 | 33 | 0 | 0 | 1.5E-02 | 17 |
| GO:0042254 | ribosome biogenesis | BP | 1.8E-05 | 12 | 0 | 0 | 1.0E-16 | 24 |
| GO:0008092 | cytoskeletal protein binding | MF | 2.0E-05 | 20 | 0 | 0 | 2.5E-13 | 36 |
| GO:0019748 | secondary metabolic process | BP | 2.4E-05 | 36 | 0 | 0 | 1.3E-02 | 18 |
| GO:0007034 | vacuolar transport | BP | 2.9E-05 | 27 | 3.2E-02 | 9 | 1.3E-02 | 16 |
| GO:0051604 | protein maturation | BP | 2.9E-05 | 29 | 0 | 0 | 2.9E-03 | 20 |
| GO:0008233 | peptidase activity | MF | 2.9E-05 | 20 | 1.2E-03 | 9 | 1.5E-03 | 16 |
| GO:0016874 | ligase activity | MF | 4.4E-05 | 24 | 0 | 0 | 2.4E-05 | 26 |
| GO:0050877 | nervous system process | BP | 4.6E-05 | 29 | 0 | 0 | 2.5E-06 | 35 |
| GO:0051301 | cell division | BP | 4.7E-05 | 19 | 2.8E-02 | 6 | 2.7E-14 | 36 |
| GO:0000902 | cell morphogenesis | BP | 5.2E-05 | 18 | 0 | 0 | 1.0E-13 | 32 |
| GO:0003723 | RNA binding | MF | 7.4E-05 | 15 | 3.1E-05 | 9 | 4.7E-10 | 24 |
| GO:0003729 | mRNA binding | MF | 8.5E-05 | 13 | 0 | 0 | 9.0E-07 | 17 |
| GO:0005730 | nucleolus | CC | 1.2E-04 | 12 | 0 | 0 | 1.3E-12 | 22 |
| GO:0040007 | growth | BP | 1.5E-04 | 15 | 0 | 0 | 1.7E-14 | 29 |
| GO:0016853 | isomerase activity | MF | 1.7E-04 | 22 | 0 | 0 | 3.9E-09 | 37 |
| GO:0007005 | mitochondrion organization | BP | 1.7E-04 | 22 | 0 | 0 | 2.6E-05 | 25 |
| GO:0006914 | autophagy | BP | 2.1E-04 | 21 | 0 | 0 | 2.0E-05 | 26 |
| GO:0005929 | cilium | CC | 4.3E-04 | 22 | 0 | 0 | 4.0E-18 | 59 |
| GO:0005777 | peroxisome | CC | 5.0E-04 | 24 | 1.1E-02 | 12 | 7.4E-03 | 18 |
| GO:0030234 | enzyme regulator activity | MF | 5.1E-04 | 16 | 3.4E-03 | 9 | 1.4E-11 | 34 |
| GO:0005815 | microtubule organizing center | CC | 5.1E-04 | 22 | 0 | 0 | 4.9E-09 | 42 |
| GO:0016791 | phosphatase activity | MF | 5.5E-04 | 19 | 3.1E-04 | 13 | 6.0E-03 | 16 |
| GO:0019843 | rRNA binding | MF | 6.8E-04 | 16 | 0 | 0 | 2.5E-06 | 25 |
| GO:0044403 | symbiont process | BP | 7.2E-04 | 20 | 0 | 0 | 5.8E-07 | 32 |
| GO:0030198 | extracellular matrix organization | BP | 7.4E-04 | 30 | 0 | 0 | 1.3E-02 | 20 |
| GO:0007155 | cell adhesion | BP | 7.7E-04 | 22 | 0 | 0 | 8.1E-07 | 37 |
| GO:0006913 | nucleocytoplasmic transport | BP | 8.0E-04 | 16 | 0 | 0 | 4.3E-09 | 31 |
| GO:0022607 | cellular component assembly | BP | 8.4E-04 | 15 | 1.8E-02 | 7 | 5.5E-09 | 27 |
| GO:0005615 | extracellular space | CC | 9.3E-04 | 18 | 1.2E-03 | 12 | 2.1E-02 | 12 |
| GO:0016757 | transferase activity, transferring glycosyl groups | MF | 1.1E-03 | 24 | 0 | 0 | 3.2E-02 | 14 |
| GO:0006457 | protein folding | BP | 1.2E-03 | 20 | 0 | 0 | 4.2E-06 | 31 |
| GO:0016765 | transferase activity, transferring alkyl or aryl (other than methyl) groups | MF | 1.2E-03 | 26 | 0 | 0 | 2.1E-02 | 16 |
| GO:0071554 | cell wall organization or biogenesis | BP | 1.4E-03 | 19 | 3.8E-06 | 19 | 0 | 0 |
| GO:0008289 | lipid binding | MF | 1.9E-03 | 18 | 0 | 0 | 3.7E-04 | 22 |
| GO:0030705 | cytoskeleton-dependent intracellular transport | BP | 2.3E-03 | 29 | 0 | 0 | 5.5E-06 | 50 |
| GO:0005198 | structural molecule activity | MF | 2.3E-03 | 24 | 0 | 0 | 1.9E-04 | 32 |
| GO:0005635 | nuclear envelope | CC | 3.2E-03 | 19 | 0 | 0 | 1.2E-08 | 40 |
| GO:0003677 | DNA binding | MF | 5.7E-03 | 12 | 3.2E-02 | 6 | 2.3E-19 | 38 |
| GO:0006399 | tRNA metabolic process | BP | 6.3E-03 | 15 | 0 | 0 | 5.9E-10 | 36 |
| GO:0022618 | ribonucleoprotein complex assembly | BP | 6.9E-03 | 9 | 0 | 0 | 2.4E-09 | 24 |
| GO:0008283 | cell proliferation | BP | 6.9E-03 | 13 | 5.0E-03 | 8 | 2.0E-11 | 34 |
| GO:0005622 | intracellular | CC | 7.4E-03 | 21 | 0 | 0 | 1.1E-06 | 47 |
| GO:0003700 | DNA-binding transcription factor activity | MF | 7.4E-03 | 17 | 0 | 0 | 3.3E-03 | 20 |
| GO:0140014 | mitotic nuclear division | BP | 7.8E-03 | 20 | 0 | 0 | 7.9E-04 | 28 |
| GO:0008135 | translation factor activity, RNA binding | MF | 8.0E-03 | 19 | 4.5E-02 | 13 | 2.5E-04 | 31 |
| GO:0007049 | cell cycle | BP | 8.1E-03 | 14 | 0 | 0 | 4.6E-12 | 39 |
| GO:0071941 | nitrogen cycle metabolic process | BP | 8.5E-03 | 40 | 0 | 0 | 4.5E-02 | 20 |
| GO:0007009 | plasma membrane organization | BP | 8.9E-03 | 50 | 0 | 0 | 4.7E-02 | 25 |
| GO:0048870 | cell motility | BP | 1.0E-02 | 13 | 0 | 0 | 1.3E-20 | 49 |
| GO:0005737 | cytoplasm | CC | 1.5E-02 | 10 | 8.7E-04 | 6 | 4.7E-39 | 37 |
| GO:0000278 | mitotic cell cycle | BP | 1.5E-02 | 11 | 2.4E-03 | 8 | 2.2E-19 | 40 |
| GO:0043226 | organelle | CC | 2.1E-02 | 17 | 8.7E-04 | 22 | 2.2E-02 | 17 |
| GO:0040011 | locomotion | BP | 2.5E-02 | 14 | 0 | 0 | 8.3E-10 | 57 |
| GO:0016810 | hydrolase activity, acting on carbon-nitrogen (but not peptide) bonds | MF | 2.5E-02 | 18 | 0 | 0 | 0 | 0 |
| GO:0016746 | transferase activity, transferring acyl groups | MF | 3.0E-02 | 14 | 1.8E-02 | 14 | 2.8E-07 | 48 |
| GO:0006259 | DNA metabolic process | BP | 3.1E-02 | 9 | 0 | 0 | 4.6E-28 | 38 |
| GO:0051082 | unfolded protein binding | MF | 3.5E-02 | 15 | 0 | 0 | 2.8E-04 | 38 |
| GO:0005654 | nucleoplasm | CC | 3.6E-02 | 8 | 0 | 0 | 1.1E-38 | 38 |
| GO:0003924 | GTPase activity | MF | 3.6E-02 | 11 | 0 | 0 | 2.9E-08 | 40 |
| GO:0000229 | cytoplasmic chromosome | CC | 4.2E-02 | 33 | 6.3E-03 | 67 | 0 | 0 |
| GO:0004518 | nuclease activity | MF | 4.5E-02 | 10 | 1.7E-02 | 9 | 7.4E-09 | 34 |
| GO:0005811 | lipid droplet | CC | 4.5E-02 | 20 | 0 | 0 | 5.0E-04 | 60 |
| GO:0032196 | transposition | BP | 4.9E-02 | 25 | 0 | 0 | 4.3E-02 | 25 |
| GO:0031012 | extracellular matrix | CC | 5.0E-02 | 20 | 0 | 0 | 7.6E-03 | 40 |
| GO:0016798 | hydrolase activity, acting on glycosyl bonds | MF | 0 | 0 | 4.9E-03 | 13 | 3.4E-02 | 13 |
| GO:0051276 | chromosome organization | BP | 0 | 0 | 4.9E-02 | 5 | 6.5E-24 | 38 |
| GO:0000228 | nuclear chromosome | CC | 0 | 0 | 0 | 0 | 7.8E-14 | 39 |
| GO:0005694 | chromosome | CC | 0 | 0 | 0 | 0 | 1.6E-10 | 45 |
| GO:0006397 | mRNA processing | BP | 0 | 0 | 0 | 0 | 2.2E-08 | 30 |
| GO:0007059 | chromosome segregation | BP | 0 | 0 | 0 | 0 | 7.5E-08 | 38 |
| GO:0008134 | transcription factor binding | MF | 0 | 0 | 0 | 0 | 1.3E-07 | 50 |
| GO:0008168 | methyltransferase activity | MF | 0 | 0 | 0 | 0 | 2.3E-07 | 36 |
| GO:0030674 | protein binding, bridging | MF | 0 | 0 | 0 | 0 | 9.6E-07 | 41 |
| GO:0043473 | pigmentation | BP | 0 | 0 | 0 | 0 | 1.5E-06 | 71 |
| GO:0004386 | helicase activity | MF | 0 | 0 | 0 | 0 | 9.0E-05 | 29 |
| GO:0016779 | nucleotidyltransferase activity | MF | 0 | 0 | 0 | 0 | 1.9E-04 | 28 |
| GO:0034330 | cell junction organization | BP | 0 | 0 | 0 | 0 | 8.5E-04 | 27 |
| GO:0042393 | histone binding | MF | 0 | 0 | 0 | 0 | 4.3E-03 | 27 |

### Supplementary figures

**Figure S1.**
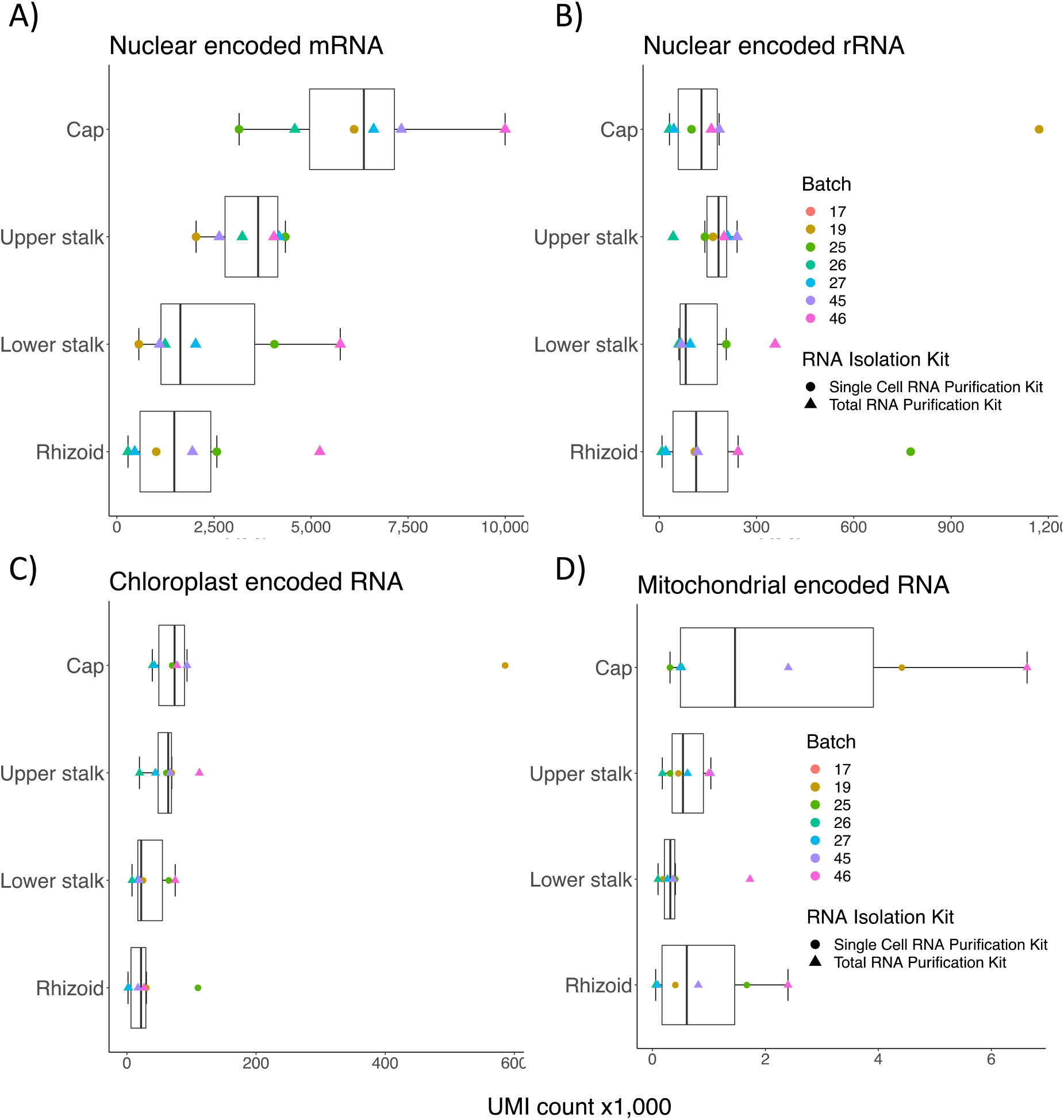
Boxplots showing the summarized gene expression levels (total UMI counts) of **A)** nuclear encoded mRNA, **B)** nuclear encoded rRNA, **C)** chloroplast encoded RNA and **D)** mitochondrial encoded RNA

**Figure S2.**
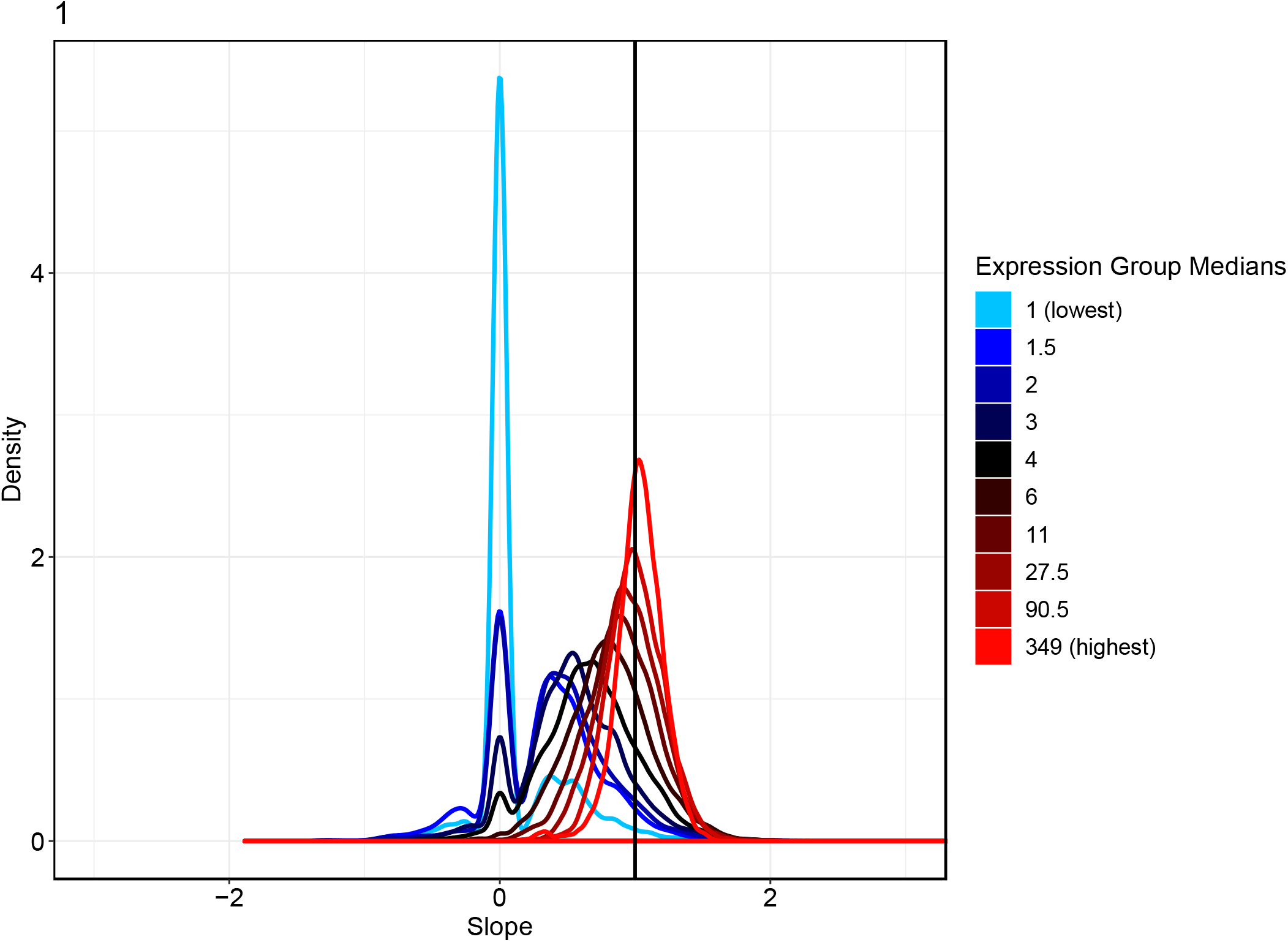
Density plot generated with the sc-norm package in R, displaying the relationship between sequencing depth and gene counts in our samples. Genes are grouped into 10 groups based on gene expression (shown in colors). We see that for the highest expressed genes, the slope of the relationship between expression and sequencing depth is close to 1, justifying the use of normalization based on sequencing depth (as in the DESeq2 R package).

